# The adhesion GPCRs CELSR1-3 and LPHN3 engage G proteins via distinct activation mechanisms

**DOI:** 10.1101/2023.04.02.535287

**Authors:** Duy Lan Huong Bui, Andrew Roach, Jingxian Li, Sumit J. Bandekar, Elizabeth Orput, Ritika Raghavan, Demet Araç, Richard Sando

**Affiliations:** Department of Pharmacology, Vanderbilt Brain Institute, Vanderbilt University, Nashville, TN 37240 USA; Department of Biochemistry and Molecular Biology, The University of Chicago, Chicago, IL, 60637, USA

## Abstract

Adhesion GPCRs (aGPCRs) are a large GPCR class that direct diverse fundamental biological processes. One prominent mechanism for aGPCR agonism involves autoproteolytic cleavage, which generates an activating, membrane-proximal tethered agonist (TA). How universal this mechanism is for all aGPCRs is unclear. Here, we investigate G protein induction principles of aGPCRs using mammalian LPHN3 and CELSR1-3, members of two aGPCR families conserved from invertebrates to vertebrates. LPHNs and CELSRs mediate fundamental aspects of brain development, yet CELSR signaling mechanisms are unknown. We found that CELSR1 and CELSR3 are cleavage-deficient, while CELSR2 is efficiently cleaved. Despite differential autoproteolysis, CELSR1-3 all engage GαS, and CELSR1 or CELSR3 TA point mutants retain GαS coupling activity. CELSR2 autoproteolysis enhances GαS coupling, yet acute TA exposure alone is insufficient. These studies support that aGPCRs signal via multiple paradigms and provide insights into CELSR biological function.

## Introduction

G protein-coupled receptors (GPCRs) are the largest and one of the most evolutionarily diverse membrane protein superfamilies in mammals^1^. GPCRs direct multitudes of fundamental biological processes by linking extracellular stimuli to intracellular responses. Adhesion-class GPCRs (aGPCRs) are a large GPCR class, with 33 members in humans, yet their biological functions and signal transduction mechanisms remain incompletely understood^2–5^. A major obstacle is the lack of a universal understanding of how the aGPCR class is agonised and how they control signal transduction cascades^6, 7^. Virtually all aGPCRs have a GAIN (GPCR autoproteolysis-inducing) domain, which harbors a GPS (GPCR proteolysis site) capable of autoproteolysis, thereby generating an extracellular N-terminal fragment (NTF) and 7-transmembrane GPCR C-terminal fragment (CTF)^8^. The most extensively studied activation mechanisms for aGPCRs is either removal or non-covalent attachment of their extracellular region following autoproteolysis, allowing a membrane proximal tethered agonist (TA) peptide, also known as the *Stachel* peptide, to agonize the 7-transmembrane GPCR and increase G protein signaling^9, 10^. How universal these TA exposure-dependent or-independent mechanisms of aGPCR activation are across all aGPCRs remains unclear.

Recent structural studies have defined the TA-mediated activation mechanism of several aGPCRs, including Latrophilin-3 (LPHN3/ADGRL3), ADGRG1, ADGRG2, ADGRG4, ADGRG5, ADGRD1, and ADGRF1, thereby providing crucial insights into TA-dependent activation^11–15^. Many aGPCRs have displayed important TA-dependent functions; for example, ADGRG6 and ADGRD1 in zebrafish studies^16^. Interestingly, cleavage-independent signaling modes and functions have been previously reported, including evidence for TA exposure in the absence of cleavage^17^. Certain aGPCRs may contain a cleavage-deficient GAIN domain, as previously shown for human BAI3 (brain angiogenesis inhibitor-3)^8^. Moreover, ADGRG5 is not efficiently autoproteolyzed yet can still become activated via its TA^18^. GPS motif presence but not cleavage of *C. elegans* LAT-1 is essential for rescuing developmental deficits in lat-1 mutants^19^. A LPHN3 autoproteolysis mutant was capable of rescuing deficits in synapse formation in LPHN3-deficient neurons^20^. In signal transduction studies, autoproteolysis has been shown to be dispensable for ADGRD1 coupling to Gαs^21^. Autoproteolysis-independent G protein mediated signaling has also been reported for GPR56/ADGRG1. Namely, synthetic ligands that bind to the extracellular region of ADGRG1 were capable of modulating signaling in ADGRG1 mutants that diminish autoproteolysis^22^. Also, an ADGRG1 mutant completely lacking a TA exhibited a subset of preserved signaling capabilities, while a BAI1 mutant lacking a TA displayed robust and fully preserved signaling capabilities^23^. Collectively, increasing evidence supports that several distinct modes of signaling including both cleavage-dependent and-independent paradigms exist for aGPCRs. How certain aGPCRs engage these distinct modes to orchestrate specific biological functions remains poorly understood.

aGPCRs have emerging roles in critical aspects of diverse biological processes. Latrophilins and CELSRs (cadherin EGF LAG-repeat 7-transmembrane receptors) are aGPCRs conserved from invertebrate to vertebrates, suggesting they underlie evolutionarily ancient biological functions^24, 25^. The mammalian Latrophilins (LPHN1-3/ADGRL1-3) and their extracellular interaction partners Teneurins and FLRTs (fibronectin leucine-rich repeat transmembrane proteins) function in fundamental aspects of synapse formation and circuit wiring specificity^20, 26–29^. LPHNs have also displayed other important roles during early brain development, including neuronal migration^30^. As GPCRs, LPHNs and their invertebrate orthologs have been shown to interact with numerous G protein-dependent pathways. *C. elegans* LAT-1 has been shown to interact with a Gαs pathway^31^, while *Drosophila* CIRL (Calcium-independent receptor for latrotoxin) Gαi^32^. In mammals, LPHN1 has displayed interaction with Gαs^33^, Gαi^33, 34^, Gαo, and Gαq pathways^35^. LPHN2 is currently less examined, but has exhibited coupling to Gαs^36, 37^, and Gα12/13 (Pederick *et al*., *BioRxiv*). LPHN3 has shown several G protein interactions, including Gαs^15, 37, 38^, Gαi^15, 34, 38^, Gαq^15, 38, 39^ and Gα12/13^11, 15, 38, 39^. The G protein dependent engagement mechanisms and signaling pathways of CELSRs remain unexplored despite their crucial roles in neurodevelopment, although a connection to the RhoA/ROCK (Rho-associated protein kinase) pathway via the planar cell polarity pathway has been illustrated^40^.

Functionally, initial studies found that the *Drosophila* CELSR ortholog starry night (Flamingo) was essential for tissue polarity via Frizzled^41–43^. Seminal neurobiological studies found that Flamingo mutants exhibited disruptions in *Drosophila* visual system topographic maps, suggesting a role in neuronal target selection during circuit assembly^44, 45^. The single *C. elegans* CELSR ortholog, Flamingo/cFMI-1, underlies a crucial role in axonal pathfinding and neuronal development^46, 47^. cFMI-1 mutants exhibited substantial deficits in GABAergic motorneuron axonal pathfinding, synapse density, and accumulation of synaptic vesicles at non-synaptic regions^48^. cFMI has been shown to genetically interact with the Wnt signaling pathway for proper neurite targeting functions^49, 50^. In the *Drosophila* nervous system, Flamingo/starry night functions in limiting dendrite extension and promoting axon extension from sensory and motor systems^51^. Important studies showed that *Drosophila* Flamingo functions as a short-range signal to influence growth cone choice of correct postsynaptic partners^52^, supporting a role in circuit wiring during development. Interestingly, the Flamingo mutant phenotype was partially rescued by a mutant lacking most of the Flamingo extracellular region, suggesting a critical role of the C-terminal 7-transmembrane GPCR region^53^. Despite these important previous studies, the biological functions of mammalian CELSRs and how they signal as aGPCRs remain unclear.

Here, we investigate autoproteolytic cleavage and G protein coupling mechanisms of the evolutionarily conserved aGPCRs CELSR1-3. We combine aGPCR protein engineering approaches to enable acute TA exposure^39, 54^ with a panel of BRET2 (bioluminescence resonance energy transfer) sensors for the GPCR “transducerome” of 14 different Gαβγ combinations^55^ to assess TA exposure-dependent and autoproteolysis-dependent modes of G protein coupling. Furthermore, we determined the differential spatial expression profiles of *Celsr1-3* during postnatal neural circuit assembly. Our results show that different mammalian CELSR isoforms induce G proteins in autoproteolysis-dependent or-independent manners and display partially non-overlapping spatial expression patterns during neurodevelopment. These studies reveal new insights into how aGPCRs orchestrate intracellular signal transduction.

## Results

### Mammalian CELSR1-3 exhibit distinct autoproteolytic cleavage profiles

Efficient autoproteolysis in the GAIN domain of LPHNs and several other aGPCRs has been well-established. However, it remains unclear if CELSRs display efficient autoproteolysis. We first set out to assess the autoproteolytic cleavage events present in *Mus musculus* (*Mm*) full-length CELSR1-3. Multiple sequence alignments showed that *Mus musculus* and *Homo sapiens* CELSR1 and CELSR3 display an unusual GAIN domain compared to other aGPCRs including LPHN1-3, with the catalytic threonine/serine cleavage residue that is part of the typical HL/T or HM/T (/ denotes cleavage site) autoproteolysis sequence replaced with an alanine or glycine, respectively (Figure 1A and B). *Mus musculus* and *Homo sapiens* CELSR2 exhibit a GAIN domain containing the catalytic threonine. The ancestral CELSR ortholog in *C. elegans* (Flamingo/FMI-1) contains a glycine residue at the typical autoproteolytic catalytic region and is more comparable to mammalian CELSR3 (Figure 1B).

**Figure 1:**
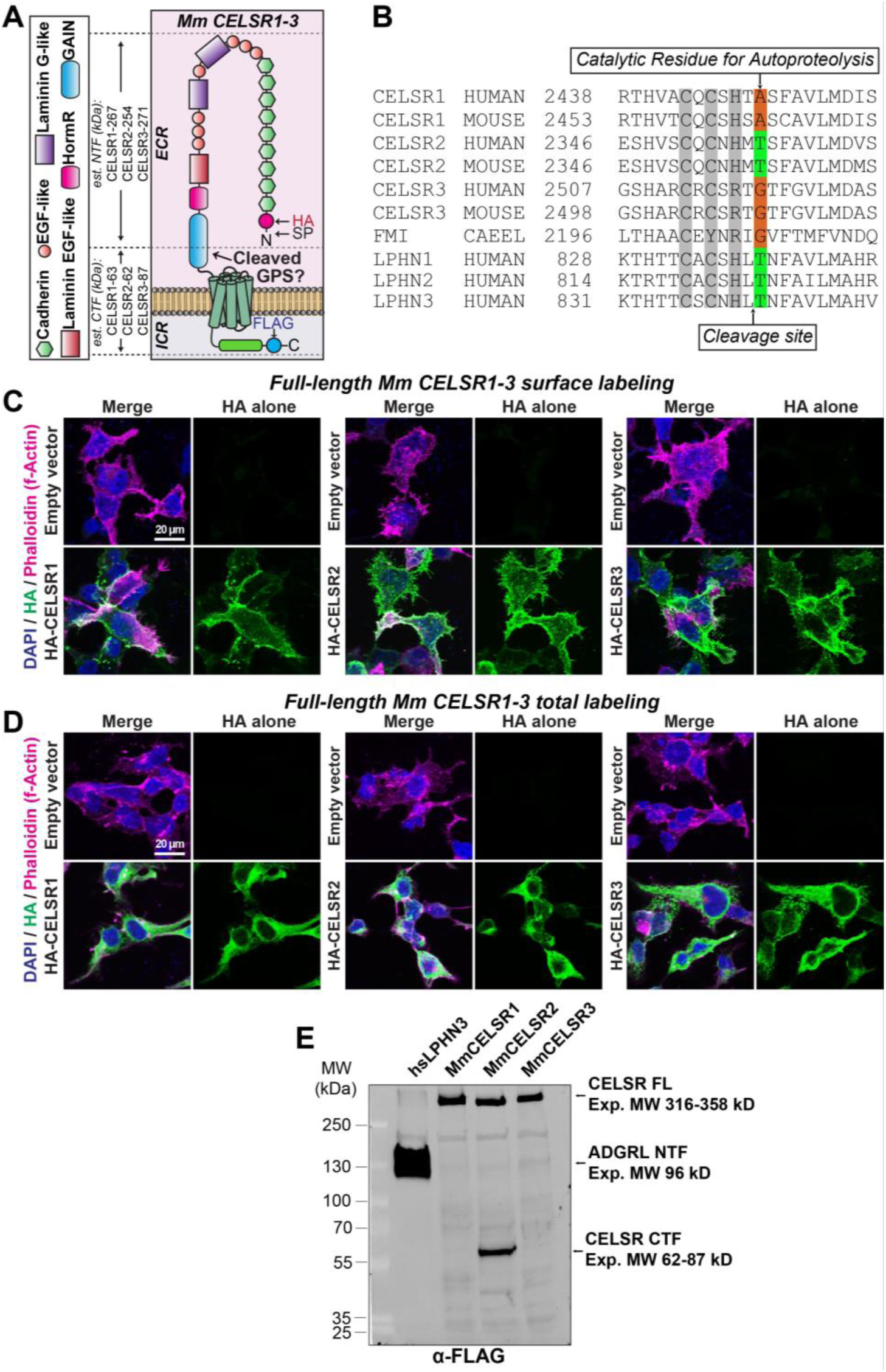
*Mm* CELSR1-3 display distinct autoproteolytic cleavage profiles. **A**, schematic diagram of full-length *Mus musculus* (*Mm*) CELSR1-3 depicting locations of N-terminal HA and C-terminal FLAG tags. The estimated molecular weight of the N-terminal fragment (NTF) and C-terminal fragment (CTF) that would result from autoproteolytic cleavage is shown on the left. ECR – extracellular region; ICR – intracellular region; EGF – epidermal growth factor; HormR – Hormone receptor domain; GAIN – G protein-coupled receptor autoproteolysis-inducing domain; SP – signal peptide; GPS – G protein-coupled receptor proteolysis site. Note that CELSR2 is predicted to contain three membrane-proximal EGF-like domains while CELSR1/3 four (three shown in Figure). **B,** multiple sequence alignment of the GAIN region of human, mouse, and *C. elegans* (CAEEL) CELSR (FMI-1) compared to human Lphn1-3 (Adgrl1-3). The autoproteolytic cleavage site is shown with an arrow on the bottom of the alignment, while the residue critical for autoproteolysis is depicted on the top of the alignment. **C,** surface expression and localization of full-length *Mm* CELSR1-3 in HEK293T cells. Cells were transfected with indicated full-length HA-tagged *Mm* CELSR constructs or empty vector, and immunolabeled for surface HA in unpermeabilized conditions, followed by Phalloidin (f-Actin) and DAPI as internal controls. **D,** total expression of full-length *Mm* CELSR1-3. Similar to C, except HEK293T cells were permeabilized and labeled for total HA together with Phalloidin and DAPI. **E,** full-length *Mm* CELSR1-3 cleavage assays in HEK293T cells. Constructs contained an N-terminal HA following a preprotrypsin signal peptide, and C-terminal FLAG tag. Full-length *Mm* CELSR2 generates a cleaved C-terminal product at the expected size following autoproteolytic cleavage, while CELSR1/3 is uncleaved under similar conditions. Human LPHN3, which exhibits efficient autoproteolysis, was used as a positive control. Expected sizes (kDa): *Mm* CELSR1 full-length (FL) – 330; NTF – 267; CTF – 63; *Mm* CELSR2 FL – 316; NTF – 254; CTF – 62; *Mm* CELSR3 FL – 359; NTF – 271; CTF – 87; *hs* LPHN3 FL – 166; NTF – 96; CTF – 70. Note that for the *hs* LPHN3 positive control, the FLAG tag was present on the N-terminal fragment. Presence of two bands in *hs* LPHN3 likely due to glycosylation (Araç *et al*., 2012). See Figure S1 and S2 for additional full-length *Mm* CELSR1-3 autoproteolysis data.

We began by cloning full-length *Mm* CELSR1-3, fusing an N-terminal HA tag and C-terminal FLAG tag to allow detection of the putative N-and C-terminal fragments, respectively (Figure 1A). We tested surface vs. total expression in HEK293T cells via immunocytochemistry and found that full-length tagged *Mm* CELSR1-3 were efficiently overexpressed and localized to the surface of HEK293T cells (Figure 1C and D). While these high-magnification images comparing surface to total immunolabeling illustrate the cell surface localization of our tagged full-length constructs, low-magnification images show the distribution of intensities across the cell population (Figure S1A and B). We subsequently used these constructs to assay autoproteolytic cleavage of full-length *Mm* CELSR1-3.

We used our tagged full-length *Mm* CELSR1-3 constructs to monitor the putative N-or C-terminal fragments resulting from potential autoproteolytic cleavage via HA or FLAG immunoblotting, respectively (Figure 1E and Figure S1C and D). Our experiments included tagged human LPHN3, which is established to be effectively cleaved, as a positive control. When transfected into HEK293T cells, we observed HA and FLAG products corresponding to the predicted sizes of cleaved N-and C-terminal fragments of CELSR2 (Figure 1E and Figure S1C and D). However, N-and C-terminal tagged products for CELSR1 and CELSR3 corresponded predominately to uncleaved, full-length receptors (Figure 1E and Figure S1C and D). Thus, consistent with predictions based on sequence alignments (Figure 1B), full-length *Mm* CELSR1 and CELSR3 are cleavage-deficient in their GAIN domain while full-length CELSR2 displays efficient autoproteolysis.

More extensive multiple sequence alignments of the GAIN region of numerous mammalian and non-mammalian CELSR1 orthologs revealed that the important catalytic autoproteolytic cleavage residue and/or activating phenylalanine important for autoproteolysis and TA-dependent activation have diverged in certain mammalian species (Figure S2). While many mammalian orthologs lack the typical threonine important for GAIN autoproteolysis (i.e. human, mouse, chimp, sheep, rat and cow), other mammals including cat and horse contain this residue (Figure S2A). This suggests that some mammalian CELSR1 orthologs may have diverged to be cleavage-deficient, while others may have preserved autoproteolytic cleavage activity. To examine this further, we generated a point mutation in full-length *Mm* CELSR1 to introduce this threonine (CELSR1 A2464T). Similar to our previous experiments, we fused an N-terminal HA and C-terminal FLAG tag to probe for the potential N-and C-terminal fragments, respectively, and assessed surface vs. total cell staining and distribution. Mutant CELSR1 A2464T was expressed and localized to the surface of HEK293T cells comparable to WT (Figure S2B-F). Interestingly, introduction of the threonine alone in the A2464T CELSR1 mutant failed to induce robust and efficient autoproteolysis compared to full-length WT *Mm* CELSR1, indicating that the determinants of autoproteolysis extend beyond this single residue (Figure S2G).

### Residue T2357 is critical for Mm CELSR2 autoproteolysis

Autoproteolysis of *Mm* CELSR2 generates a putative TA peptide, similar to LPHNs. A predicted residue important for autoproteolytic cleavage site of CELSR2 based on known cleaved aGPCRs and multiple sequence alignments in Figure 1B is T2357. To determine if this residue is involved, we mutated this threonine to generate two CELSR2 point mutants, T2357A and T2357G, and examined their expression and autoproteolytic cleavage compared to WT CELSR2 and empty vector (Figure 2 and Figure S3). Mutant T2357A and T2357G CELSR2 harbored an N-terminal HA and C-terminal FLAG tag similar to our WT full-length construct. Tagged CELSR2 T2357A and CELSR2 T2357G were expressed on the surface of HEK293T cells at relative levels indistinguishable from WT CELSR2, suggesting that these point mutations did not alter protein folding or cell surface localization (Figure 2A-C and Figure S3). We then tested these mutant CELSR2 forms in autoproteolysis assays and found that both point mutations diminished autoproteolytic cleavage of *Mm* CELSR2, resulting in predominately a single band via FLAG immunoblotting corresponding to the expected size of the full-length receptor (Figure 2D). Therefore, *Mm* CELSR2 requires residue T2357 for effective GAIN domain autoproteolysis.

**Figure 2:**
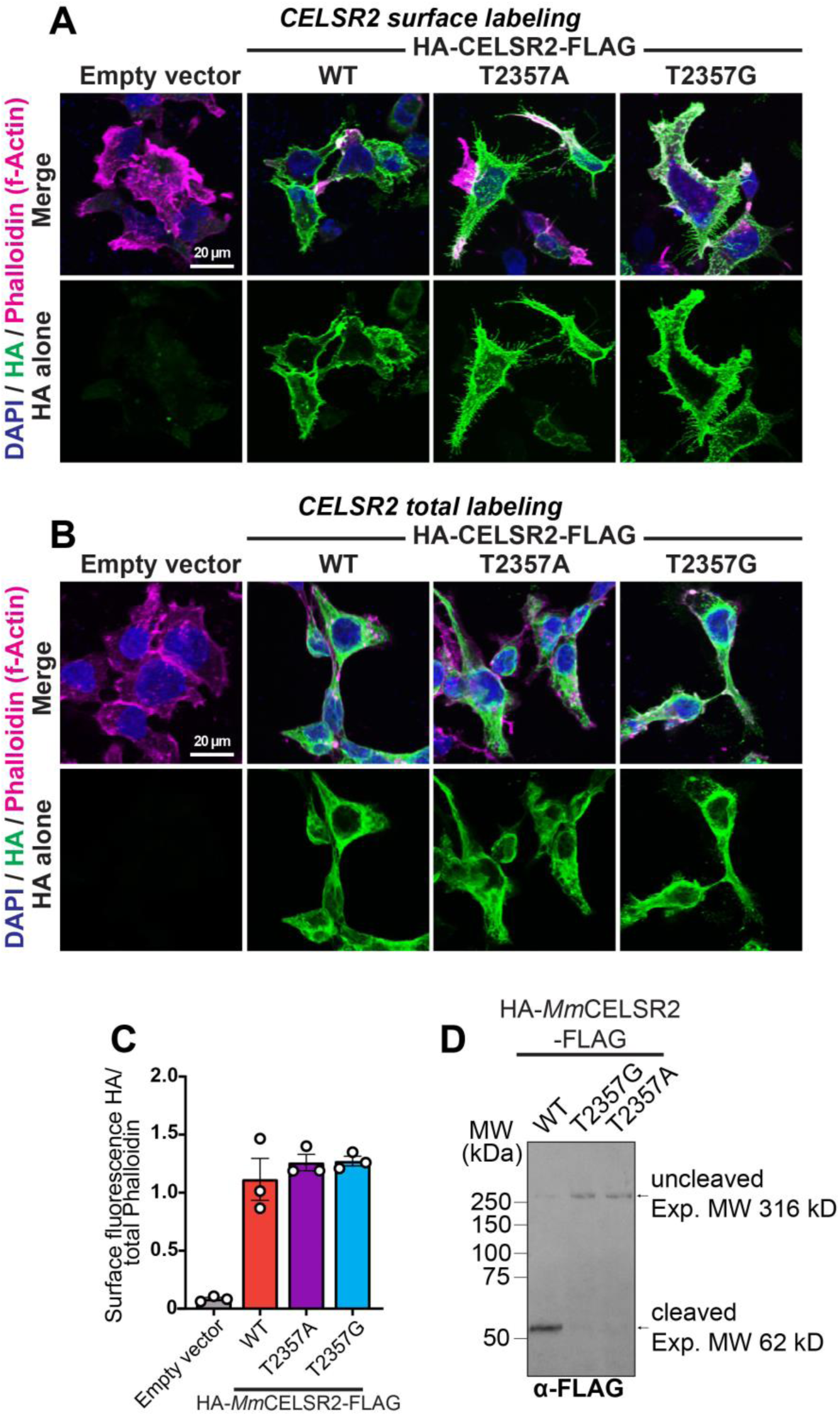
T2357 is important for *Mm* CELSR2 autoproteolysis. A, representative images of surface expression levels of full-length *Mm* CELSR2 relative to CELSR2 point mutants (T2357A and T2357G). HEK293T cells were transfected with indicated full-length HA-*Mm* CELSR2-FLAG constructs (WT or indicated point mutants) or empty vector, and immunolabeled for surface HA in unpermeabilized conditions, followed by Phalloidin (f-Actin) and DAPI (nuclei) as internal controls. **B,** total expression of indicated constructs. Similar to A, except HEK293T cells were permeabilized and labeled for total HA together with Phalloidin and DAPI. **C,** quantification of HA surface intensity relative to Phalloidin. Total cell fluorescence was measured for HA and Phalloidin channels from three independent culture replicates. **D,** *Mm* CELSR2 cleavage assays. HEK293T cells were transfected with the indicated HA-*Mm* CELSR2-FLAG constructs, and subsequently immunoblotted for the C-terminal FLAG-tagged cleavage product. Numerical data are means ± SEM from 3 independent biological replicates (depicted as open circles). See Figure S3 for additional data regarding *Mm* CELSR2 autoproteolysis.

### Evaluation of aGPCR acute TA exposure-dependent and-independent G protein coupling

Our results support that full-length *Mm* CELSR1 and CELSR3 are cleavage-deficient in mammalian cells, while CELSR2 is efficiently cleaved (Figure 1 and 2). The differential mechanisms of cleavage-dependent and-independent aGPCR:G protein coupling and G protein pathways controlled by CELSRs remain unclear. To establish a system to evaluate acute TA-dependent G protein coupling of aGPCRs, we implemented recent advances allowing for acute, thrombin-mediated exposure of the aGPCR TA peptide^39^. We combined the thrombin-mediated acute TA exposure approach with bioluminescence resonance transfer (BRET2) sensors (termed TRUPATH) that assay the “transducerome’ of 14 Gαβγ combinations (Figure 3 and Figure S4)^55^. We validated this approach using LPHN3, as in previous studies^39^. We began by replacing the large extracellular portion of *Mus musculus* LPHN3 (NCBI# NM_001359828.1) that is N-terminal of the autoproteolytic cleavage site with PAR (N-terminal domain of protease-activated receptor) (Figure 3A and B). This approach allows for overexpression of a TA-protected LPHN3, and subsequent thrombin-mediated cleavage and removal of PAR^56^, which thereby exposes the TA peptide (Figure 3B). Similar to Mathiasen *et al*., 2020, we fused the PAR sequence to LPHN3 so that thrombin-mediated cleavage in PAR at LDPR/SF would expose the critical activating phenylalanine in the endogenous LPHN3 TA, thereby generating a thrombin-exposed TA of N-SFAVLM…-C (Figure 3B).

**Figure 3:**
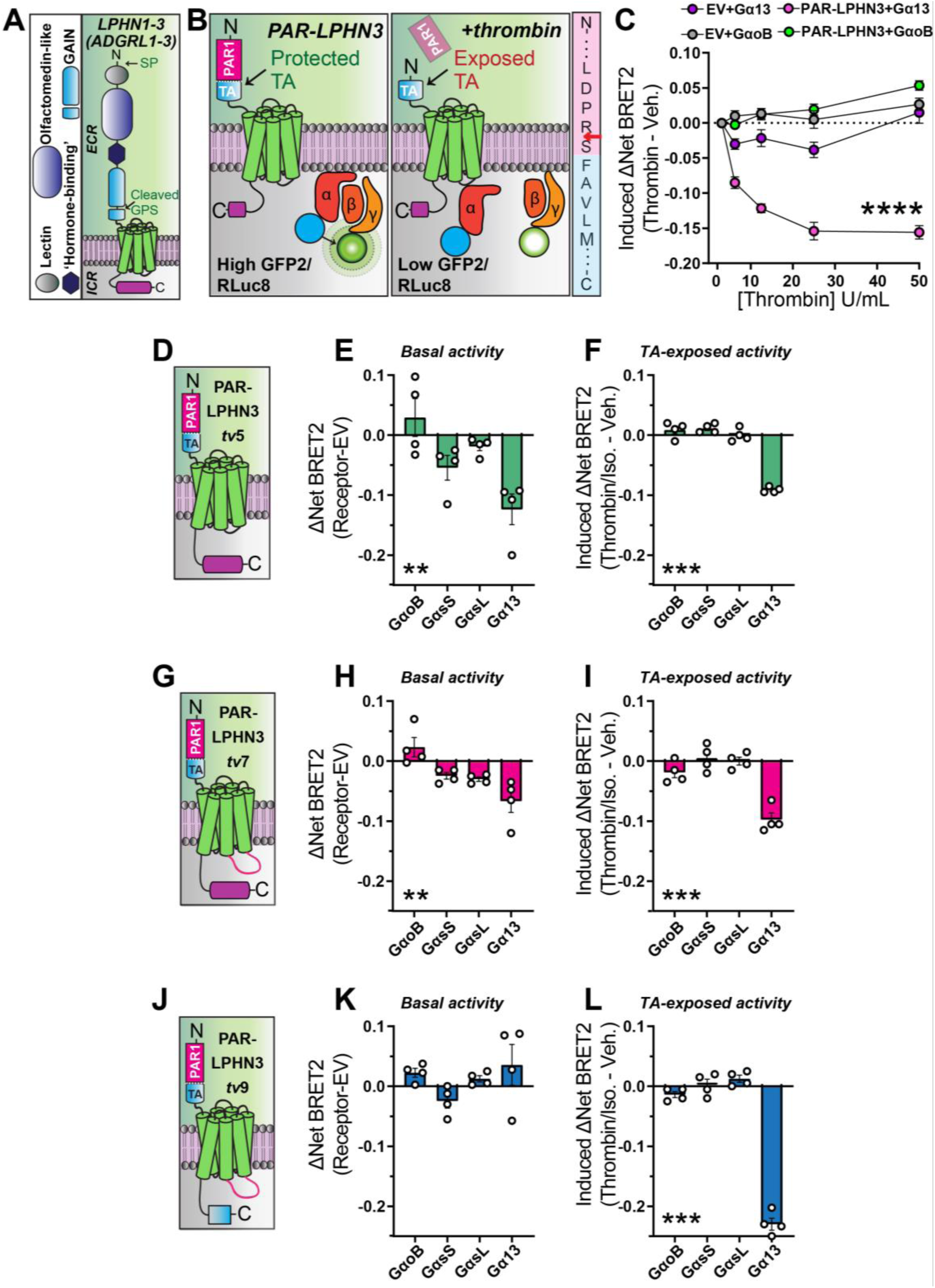
Validation of BRET2 approach to evaluate TA exposure-dependent and independent aGPCR:G protein coupling. **A,** Diagram of *Mus musculus* LPHN1-3 (ADGRL1-3) domain organization. LPHNs exhibit a cleaved GAIN domain, which exposes a self-tethered agonist ‘TA’ (i.e. *Stachel* peptide). ECR – extracellular region; ICR – intracellular region; GAIN – G-protein-coupled receptor autoproteolysis-inducing domain; SP – signal peptide; GPS – G protein-coupled receptor proteolysis site. **B,** Model of experimental approach for acute TA exposure and subsequent BRET2 measurements of G protein coupling. *Left*, replacement of the LPHN3 ECD with PAR protects the TA from inducing TA-dependent G protein induction. *Center*, thrombin-mediated cleavage and removal of PAR exposes the TA, resulting in TA-dependent activation generating a decrease in BRET2 ratio. *Right*, illustration of PAR-LPHN3 TA cleavage site. Thrombin cleaves PAR (pink sequence) following arginine (red arrow), exposing an N-terminal serine followed by the activating phenylalanine present in the LPHN3 TA sequence (light blue sequence). **C,** Thrombin-mediated LPHN3 TA exposure induces Gα13 coupling using TRUPATH G protein BRET2 biosensors. G protein KO HEK cells (GKO HEK293) were transfected with either empty vector (EV) or PAR-LPHN3 together with indicated BRET2 G protein sensor combinations. Transfected cells were subjected to increasing concentrations of thrombin for 10 minutes and BRET2 measurements were conducted. **D,** Schematic diagram of PAR-LPHN3*tv*5 (NCBI reference sequence NM_001347371.2). **E,** basal G protein coupling of PAR-LPHN3*tv*5 using the indicated set of TRUPATH BRET2 sensors. Net BRET2 signal from GKO HEK293 cells transfected with overexpressed PAR-LPHN3 variants were compared to cells transfected with empty vector which were co-transfected with the same G protein BRET2 sensors. **F,** PAR-LPHN3*tv*5 TA-exposure dependent G protein coupling. For TA-induced coupling, GKO HEK293 cells were transfected with the indicated BRET2 G protein sensor combinations and the net BRET2 signal of cells receiving 10-minute 10 U/mL thrombin treatment were compared to vehicle treated cells. **G,** Schematic diagram of PAR-LPHN3*tv*7 (NCBI reference sequence NM_001359828.1). **H,** same as E, except for PAR-LPHN3*tv*7. **I,** same as F, except for PAR-LPHN3*tv*7. **J,** Schematic diagram of PAR-LPHN3*tv*9 (NCBI reference sequence NM_001359830.1). LPHN3*tv*9 contains the same intracellular loop 3 as LPHN3*tv*7, except differs from LPHN3*tv*5 and LPHN3*tv*7 by exhibiting a distinct, shorter intracellular tail sequence. **K,** same as E, except for PAR-LPHN3*tv*9. **L,** same as F, except for PAR-LPHN3*tv*9. Numerical data are means ± SEM from four independent biological replicates (depicted as open circles in bar graphs). Statistical significance was assessed by two-way ANOVA (****, p<0.0001) in panel C and one-way ANOVA for remaining panels (**, p<0.01; ***, p<0.001). Asterisks depict ANOVA results. See Figure S4 for additional PAR-LPHN3 data.

We validated this approach with the TRUPATH sensors for Gα13 (Gα13/Gβ3/Gγ9), which was previously shown to couple LPHN3 in a TA exposure-dependent manner^39^, and GαoB (GαoB/Gβ3/Gγ8) as a negative control. BRET2 experiments were conducted in G protein KO (GKO) HEK293 cells to minimize interference with endogenous G proteins and our overexpressed receptors-of-interest (Figure 3C)^57, 58^. We transfected the same TRUPATH sensors in cells co-transfected with empty vector (EV) to control for non-specific effects of thrombin treatment. Consistent with previous studies^39^, we found thrombin dose-dependent PAR-LPHN3 coupling to Gα13. Importantly, no significant change in net BRET2 was observed in empty vector conditions treated with similar concentrations of thrombin, supporting that Gα13 coupling was specific for thrombin-mediated cleavage of LPHN3 (Figure 3C). Thus, combining the PAR-LPHN3 approach with TRUPATH sensors provides a system to assess TA-dependent G protein coupling in aGPCRs.

Multiple putative transcript variants are present in public databases for *Mm* LPHN3 in intracellular loop 3 (ICL3), which is important for G protein coupling, as well as the C-terminal intracellular tail sequence, which has been shown to modulate signal transduction pathway specificity for LPHN1^33^. We used these thrombin-dependent acute TA exposure and BRET2 approaches to determine how three putative LPHN3 sequence variants (NCBI transcript variant 5 NM_001347371.2, NCBI transcript variant 7 NM_001359828.1, and NCBI transcript variant 9 NM_001359830.1; here referred to as LPHN3*tv5*, LPHN3*tv7*, and LPHN3*tv9*, respectively) impact G protein coupling (Figure 3D-L). Given several aGPCRs have shown TA exposure-dependent and-independent activity, and LPHN3 has been reported to exhibit both basal and agonist-dependent G protein coupling^38^, we examined basal vs. TA exposure-dependent G protein coupling in parallel. As an initial control to assess potential differences in expression levels of the three PAR-LPHN3 variants tested, we immunolabeled for overexpressed PAR-LPHN3 variants containing a C-terminal HA-tag but observed no statistically significant differences (Figure S4A and B).

LPHN3*tv5* exhibits a relatively short ICL3 followed by an intracellular C-terminal tail region with a PDZ-binding motif (Figure 3D). We found that PAR-LPHN3*tv5* displayed modest basal GαsS coupling and more substantial basal Gα13 coupling in the absence of thrombin-mediated TA exposure (Figure 3E), as well as exposed TA-induced Gα13 coupling (Figure 3F). Further control experiments showed effective agonist-induced (10 µM Isoproterenol) coupling of GαsS and GαsL to β2-adrenergic receptor (β2-AR) (Figure S4C), as reported in the TRUPATH study^55^. PAR-LPHN3*tv7*, which contains a 9 amino acid insertion (NYEDNRPFI) in ICL3 compared to LPHN3*tv5* (Figure 3G), also displayed modest basal and exposure-dependent G protein coupling (Figure 3H and I). PAR-LPHN3*tv9*, which contains the same ICL3 as LPHN3*tv7* but a short C-terminal intracellular tail (Figure 3J), displayed no detectable basal Gα coupling (Figure 3K) and robust TA-dependent Gα13 coupling (Figure 3L). To control for other possibilities, such as receptor induced changes in the expression of TRUPATH sensors, we plotted the donor luminescence measurements from these experiments but found no significant differences between conditions for given sensors (Figure S4D). These experiments further validate these approaches and suggest that aGPCR transcript variants may impact G protein coupling efficacy.

Given the importance of LPHNs in mechanisms of synapse formation and brain development, we subsequently conducted RNA *in situ* hybridizations with C-terminal variant specific and pan-*Lphn3* probes at three time-points during synaptogenesis. Interestingly, the C-terminal tail present in Lphn3*tv*5 and 7 harbors a PDZ-binding motif that interacts with postsynaptic SHANK proteins^59^. We co-labeled brain sections from postnatal day 5, 10, and 21 mice for a probe detecting all *Lphn3* transcript variants (pan-*Lphn3*) together with a probe specific for the C-terminal tail sequence present in Lphn3*tv5/7* but not Lphn3*tv9* (Ct-*Lphn3*) (Figure 4 and Figure S5). Probes specific for pan-*Lphn3* as well as the Lphn3*tv5/7* C-terminal intracellular tail were robustly detected throughout the developing mouse hippocampus (Figure 4A). We quantified the intensities of each probe relative to the area occupied by DAPI and found that Lphn3 expression increased during postnatal development, particularly in the CA1 (Figure 4B and C) and CA3 (Figure 4D and E) compared to the dentate gyrus (Figure 4F and G). The pan-*Lphn3* and Ct-*Lphn3* probes did not completely co-localize in all hippocampal sub-regions analyzed (Figure S5). The small size of the ICL3 insert sequence precluded similar analyses with this sequence variant, and future studies are required to determine its biological significance *in vivo*. Altogether, these experiments support that *Lphn3* expression increases in the hippocampus during early postnatal development, and that the large C-terminal tail harboring a PDZ-binding motif is highly prevalent during this developmental period.

**Figure 4:**
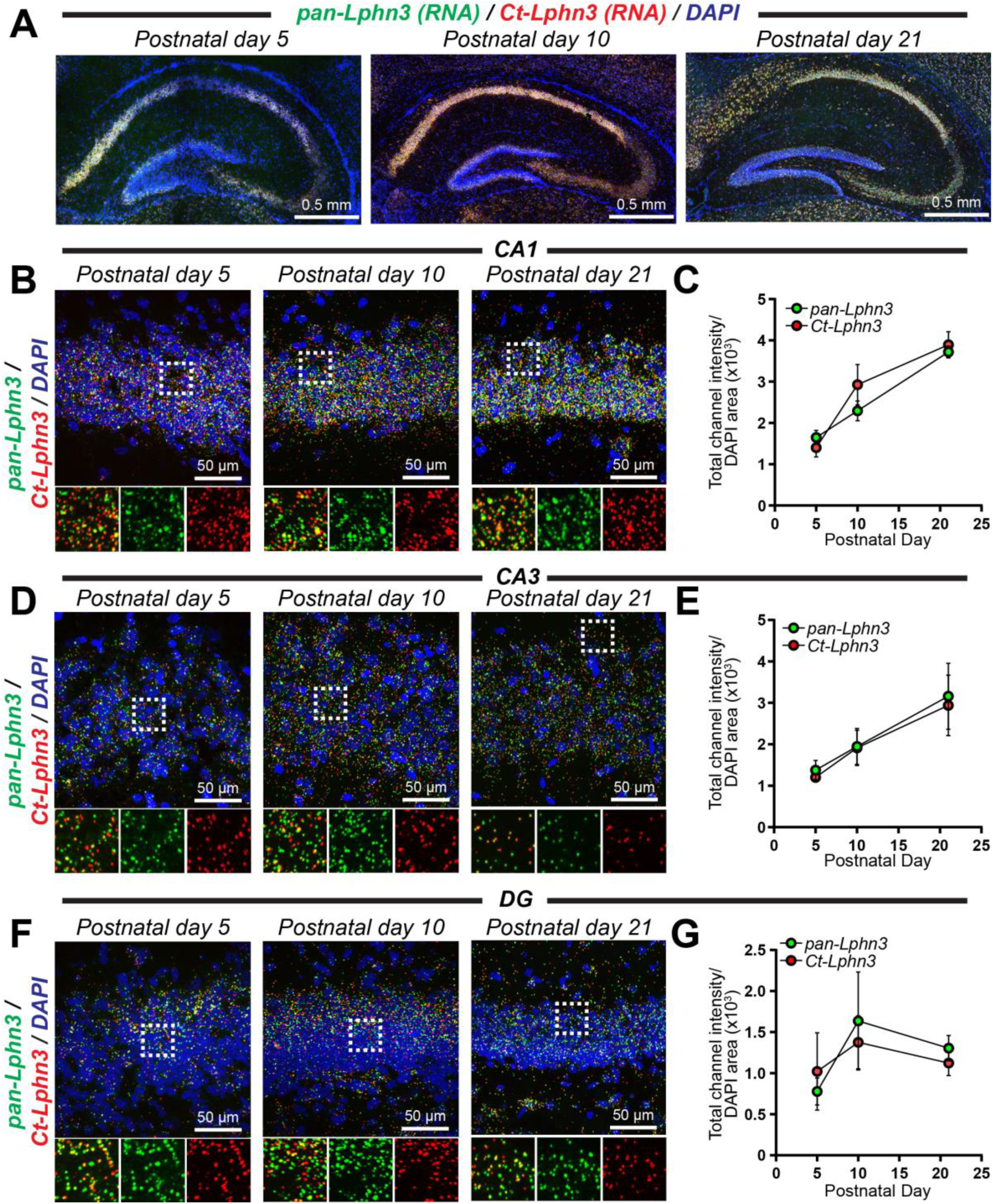
Total and splice site-specific *Lphn3* spatial expression in the developing mouse hippocampus. **A,** representative images of the postnatal day 5, 10, and 21 mouse hippocampus labeled for RNA *in situ* probes for pan-Lphn3 transcripts together with a probe specific for the C-terminal tail sequence (Ct-Lphn3) that is present in Lphn3*tv*5 and Lphn3*tv*7, but not Lphn3*tv*9. **B,** representative high-magnification 60X images of *in situs* in the hippocampal CA1 region at postnatal day 5, 10, and 21. *Top*, representative merged channel image; *bottom*, zoom-in of white boxed area from *left* with individual channels separated for pan-Lphn3 (green) and Ct-Lphn3 (red). **C,** quantifications of CA1 RNA *in situ* results. The total channel intensity of green (pan-Lphn3) or red (Ct-Lphn3) signal was compared to the area occupied by DAPI for each respective image. **D & E,** same as B and C, except for the hippocampal CA3 region. **F & G,** same as B and C, except for the hippocampal dentate gyrus (DG). Data are from 3-4 independent biological replicates. See Figure S5 for additional *Lphn3* RNA *in situ* quantifications.

### Full-length CELSR1-3 engage in G protein coupling with GαsS

While autoproteolytic cleavage of *Mm* CELSR2 will produce a putative TA peptide, our results support that full-length *Mm* CELSR1 and CELSR3 are cleavage-deficient in mammalian cells. This indicates autoproteolysis-independent modes of signaling may exist for *Mm* CELSR1 and CELSR3. To begin interrogating this, we applied the complete panel of TRUPATH BRET2 sensors to full-length CELSR1-3 to assess G protein coupling (Figure 5 and Figure S6). Given the basal Gα13 coupling we observed upon PAR-LPHN3*tv5* overexpression, we utilized this as a control within these experiments. Full-length *Mm* CELSR1 displayed GαsS coupling in these experiments (Figure 5A). Heat maps illustrate that in these experimental conditions, full-length CELSR1 coupled to GαsS within the panel of TRUPATH sensors (Figure 5B). We subsequently performed plasmid copy number titer experiments, where we increased the amount of CELSR1 plasmid transfected together with a given amount of TRUPATH GαsS sensors, balancing with empty vector to transfect the same total plasmid amount (Figure 5C). These experiments demonstrated an increasing BRET2 response with increased CELSR1 plasmid copy number, comparing to empty vector transfected cells (Figure 5C). Interestingly, similar experiments with full-length *Mm* CELSR2 and CELSR3 also revealed GαsS coupling and plasmid copy-dependent GαsS coupling (Figure 5D-I). Notably, while a direct comparison has caveats, CELSR2 induced at stronger plasmid copy-dependent GαsS response (Figure 5F) compared to CELSR1 (Figure 5C) or CELSR3 (Figure 5I). Similar to our LPHN3 BRET2 studies, we also analyzed the donor luminescence readings from BRET2 experiments but found no detectable differences between different TRUPATH sensors for a given CELSR isoform (Figure S6A-C). Altogether, these results support that full-length cleavage-deficient *Mm* CELSR1 and CELSR3 and full-length cleaved CELSR2 engage in G protein coupling with GαsS.

**Figure 5:**
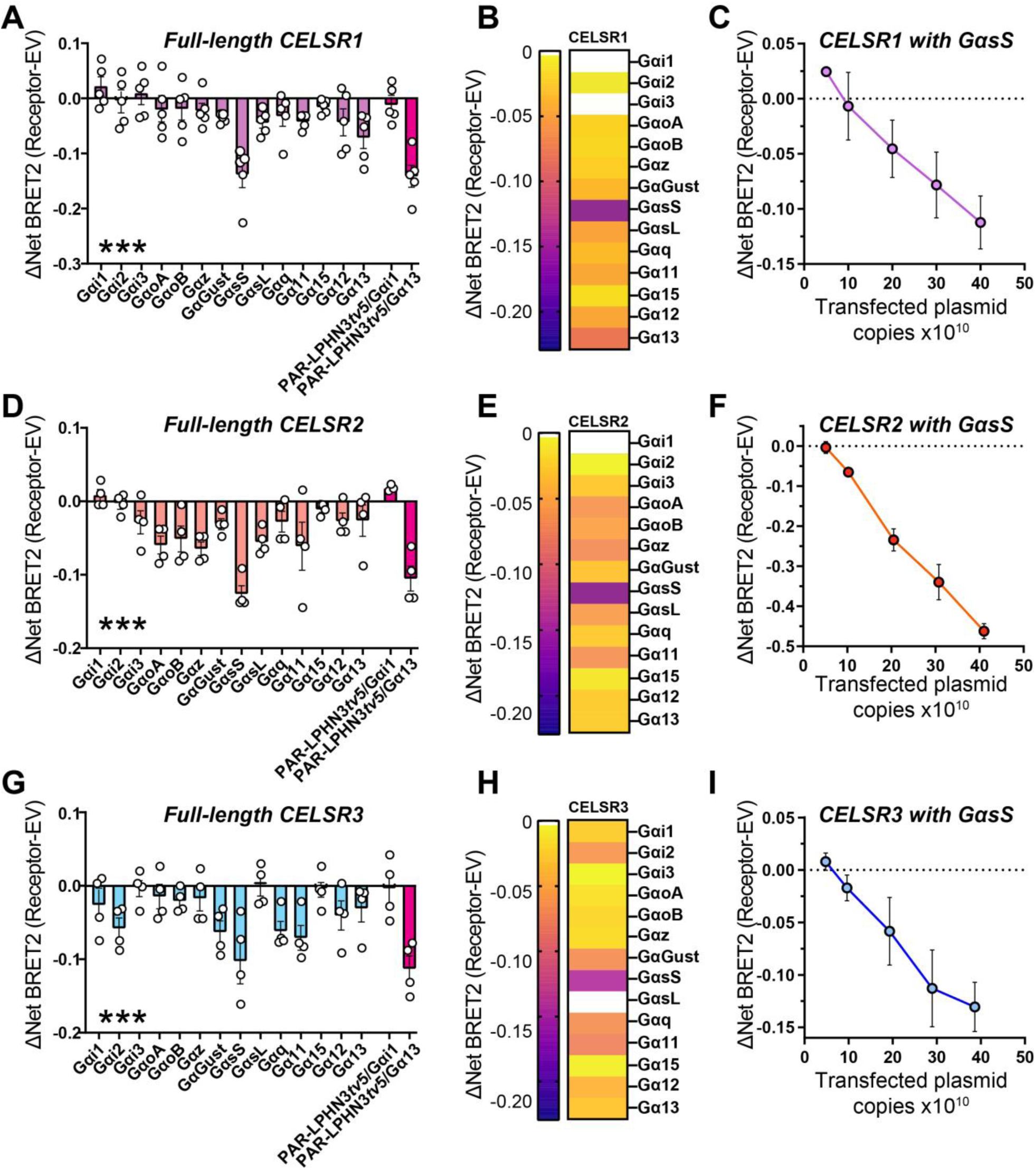
Full-length *Mm* CELSR1-3 are capable of GαsS coupling. **A,** BRET2 assays with full-length *Mm* CELSR1 and the complete panel of TRUPATH sensors in GKO HEK293 cells to test basal G protein coupling. Basal activity of PAR-LPHN3*tv*5 with Gαi1 or Gα13 were used as controls in all experiments. Data are from 5 independent biological replicates, as depicted by open circles in graphs. **B,** heat map illustrating G protein coupling of full-length *Mm* CELSR1. **C,** copy-dependent full-length *Mm* CELSR1 coupling to GαsS. GKO HEK293 cells were transfected with increasing amounts of full-length *Mm* CELSR1 and BRET2 measurements conducted compared to cells transfected with the same total amount of empty vector. Data are from 3 independent biological replicates. **D,** Full-length *Mm* CELSR2 G protein coupling via BRET2. Experiments were conducted as in panel A, except for CELSR2. Data are from 4 independent biological replicates. **E,** heat map illustrating G protein coupling of full-length *Mm* CELSR2. **F,** copy-dependent full-length *Mm* CELSR2 coupling to GαsS. Experiments were performed as in panel C and data are from 3 independent biological replicates. **G,** Full-length *Mm* CELSR3 G protein coupling via BRET2. Experiments were conducted as in panel A, except for CELSR3. Data are from 4 independent biological replicates. **H,** heat map illustrating G protein coupling of full-length *Mm* CELSR3. **I,** copy-dependent full-length *Mm* CELSR3 coupling to GαsS. Experiments were performed as in panel C and data are from 3 independent biological replicates. Numerical data are means ± SEM from 3-5 independent biological replicates (depicted as open circles in bar graphs), and as indicated in the Figures Legends. Statistical significance was assessed by one-way ANOVA (***, p<0.001). Asterisks depict one-way ANOVA results. See Figure S6 for additional CELSR BRET2 data.

### Celsr1-3 are highly abundant during neural circuit assembly and display partially non-overlapping spatial expression patterns

Despite the fundamental importance of the invertebrate CELSR orthologs Flamingo/FMI-1/starry night in circuit assembly during brain development, the functions of mammalian CELSR1-3 remain unclear. Thus, we conducted spatial expression analysis of *Mm Celsr1-3* through the developing mouse brain to obtain insights into their *in vivo* roles (Figure 6 and Figure S7). Similar to our *Lphn3* studies, we performed RNA *in situ* hybridizations for *Celsr1-3* at postnatal days 5, 10 and 21, which is a major period of synaptogenesis in the mouse brain. Interestingly, *Celsr1-3* were present in partially non-overlapping spatial patterns during postnatal development, with persistent expression through postnatal day 21 (Figure 6A and Figure S7). We then quantified the intensity of each *Celsr* isoform relative to the area occupied by DAPI in the CA1, CA3, and dentate gyrus regions during development. *Celsr2* and *Celsr3* were the predominant isoforms expressed and were largely restricted to the pyramidal cell layers of the hippocampal CA1/CA3 and granule cell layers of the dentate gyrus (Figure 6B-G). The spatial expression of *Celsr1* was distinct compared to *Celsr2* and *Celsr3*. *Celsr1* was expressed in the hippocampal region at postnatal day 5, but became increasingly restricted to the granule layer of the hippocampal dentate gyrus over development (Figure 6). While *Celsr1* was uniformly distributed throughout the dentate gyrus granule cell layer at postnatal day 5, it became selective for the subgranular zone by postnatal day 21, where cells responsible for adult neurogenesis reside (Figure 6F and G). Thus, different *Celsr* isoforms display discrete spatial expression patterns, with *Celsr1* and *Celsr2/3* being particularly distinct.

**Figure 6:**
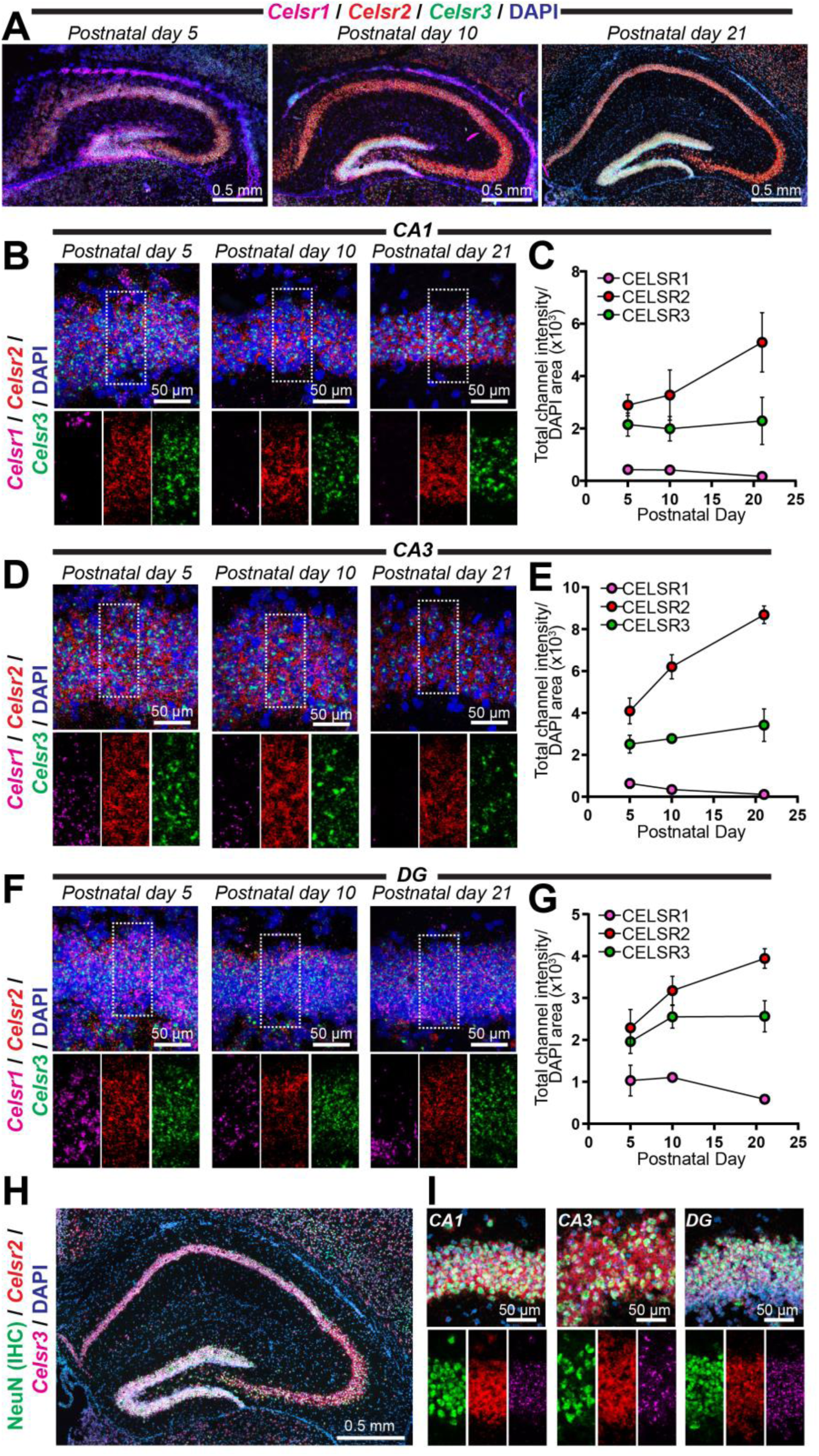
Partially overlapping *Celsr1-3* spatial expression during hippocampal circuit assembly. **A,** representative image of the postnatal day 5, 10, and 21 mouse hippocampus co-labeled with RNA *in situ* probes for *Celsr1-3* and DAPI. **B,** representative high-magnification 60X images of *in situs* in the hippocampal CA1 region at postnatal day 5, 10, and 21. *Top*, representative merged channel image; *bottom*, zoom-in highlighting the white boxed pyramidal cell layer area from *top* with individual channels separated for *Celsr1* (magenta), *Celsr2* (red), and *Celsr3* (green). **C,** quantifications of CA1 RNA *in situ* data. The total channel intensity of green (*Celsr3*), red (*Celsr2*), or far-red (*Celsr1*) signal was compared to the area occupied by DAPI for each respective image **D,** same as B, except for the hippocampal CA3 region at postnatal day 5, 10, and 21. **E,** quantifications of CA3-region RNA *in situ* data. Similar to C, except for the hippocampal CA3 at postnatal day 5, 10, and 21. **F,** same as B, except for the hippocampal dentate gyrus (DG) at postnatal day 5, 10, and 21. **G,** quantifications of RNA *in situ* data from the dentate gyrus. Similar to C, except for the hippocampal dentate gyrus at postnatal day 5, 10, and 21. **H-I,** double immunohistochemistry (IHC) and RNA *in situs* for the neuronal marker NeuN together with *Celsr2/3* RNA. **H,** representative postnatal day 21 hippocampus co-labeled for NeuN (IHC; green), *Celsr2* (RNA *in situ*; red), *Celsr3* (RNA *in situ*; magenta) and DAPI. **I,** representative high-magnification 60X image of the postnatal day 21 hippocampal CA1, CA3 and dentate gyrus from double IHC/RNA *in situ* experiments. *Top*, merged image depicting IHC for NeuN (green) together with RNA *in situs* for *Celsr2* (red), *Celsr3* (magenta), and DAPI. *Bottom*, individual channel images for the center area of top highlighting *Celsr2/3* distribution. Data are from 3-4 independent biological replicates. See Figure S7 for additional *Celsr1-3* spatial expression data.

We examined *Celsr2/3* spatial expression more globally by imaging *in situs* from sagittal sections, and found that *Celsr2* was broadly expressed, while *Celsr3* co-expressed with *Celsr2* in discrete areas including the olfactory bulb, hippocampus, and cerebellum (Figure S7). To determine if *Celsr2/3* expression in hippocampal sub-regions was enriched in neurons of the pyramidal and granule cell layers, we conducted double immunohistochemistry (IHC)/RNA *in situs* for the mature neuronal marker NeuN together with the RNA probes for *Celsr2/3* (Figure 6H and I). These experiments revealed that *Celsr2/3* expression strongly overlapped with NeuN in the CA1, CA3 and dentate gyrus (Figure 6H and I). Altogether, these studies support that *Celsr1-3* display differential autoproteolysis as well as spatial expression patterns in the developing mouse brain.

### Autoproteolysis in the context of full-length Mm CELSR2 is important for GαsS coupling

While full-length *Mm* CELSR2 exhibits GαsS coupling properties, CELSR2 autoproteolytic cleavage generates a TA peptide with potential TA exposure-dependent or-independent functions. To first address if autoproteolysis is important for CELSR2:GαsS coupling, we tested the ability of mutant T2357A CELSR2 to couple to GαsS (Figure 7A). We conducted similar plasmid copy number-dependent GαsS coupling experiments as in Figure 5 and found that the T2357A mutation significantly attenuated GαsS coupling compared to WT CELSR2 (Figure 7A). This supports that efficient *Mm* CELSR2 autoproteolysis enhances its GαsS coupling.

**Figure 7:**
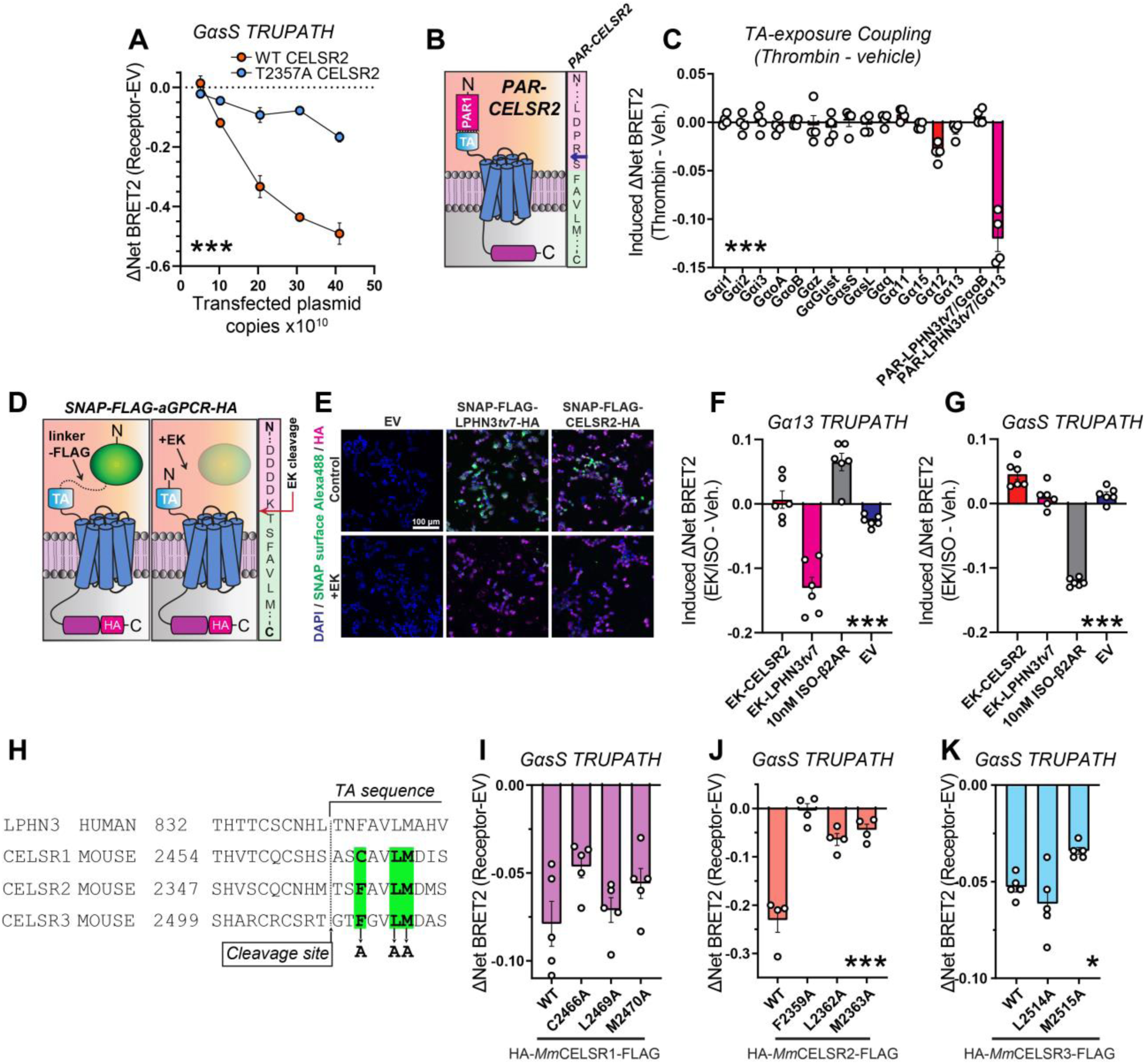
CELSRs engage distinct GαsS induction mechanisms. **A,** Plasmid dose-dependent WT CELSR2 and T2357A CELSR2 GαsS coupling. Experiments were conducted as in Figure 5, except full-length T2357A CELSR2 was analyzed in parallel to full-length WT CELSR2. **B,** *Left*, schematic diagram for PAR-CELSR2. *Right*, illustration of PAR-CELSR2 cleavage site. Thrombin cleaves PAR (pink sequence) following arginine (blue arrow), exposing an N-terminal serine followed by the TA sequence (light green sequence). **C,** G protein coupling following acute TA exposure in PAR-CELSR2 using the complete panel of TRUPATH sensors. BRET2 measurements were conducted for four independent biological replicates. TA-induced coupling of PAR-LPHN3*tv*7 with GαoB or with Gα13 were used as controls. **D,** Strategy for acute exposure of a native tethered agonist^54, 61^. A SNAP tag-linker-FLAG sequence was fused immediately upstream the native tethered agonist sequence for LPHN3*tv*7 or CELSR2. Enterokinase cleaves immediately following DDDK/ in FLAG, exposing the native TA peptide. Sequence on right shows fusion region for CELSR2 (see Methods for additional details on design). **E,** Validation of surface expression and enterokinase-mediated cleavage of SNAP tags. HEK293T cells were transfected with indicated experimental or control (EV) plasmids in 24-well plates, treated with 32 U enterokinase vs. control, and labeled for cell-impermeable SNAP ligand conjugated to Alexa Fluor 488. Cells were subsequently fixed and immunolabeled for the C-terminal HA tag as an internal control. **F,** Gα13 coupling of indicated experimental conditions. Transfected cells plated into 96-well plates were treated with 5.5U/well enterokinase (EK) or vehicle for 15 minutes, and BRET2 was subsequently measured. β2-AR treated with 10 nM ISO and EV treated with EK were used as a controls. Data are from 6 independent biological replicates. **G,** GαsS coupling in indicated conditions. Same as F, except the GαsS TRUPATH sensors were used. **H,** Multiple sequence alignment of the GAIN region of human LPHN3 and mouse CELSR1-3. Putative residues for TA-mediated agonism of CELSR1-3, which were systematically mutated to alanine to examine their contribution to GαsS induction, are highlighted in green. **I,** GαsS coupling in indicated full-length WT, C2466A, L2469A, and M2470A CELSR1 conditions. See Figure S9 for additional data on mutant expression and autoproteolysis. Data are from 5 independent biological replicates. **J,** Same as I, except for full-length WT, F2359A, L2362A, and M2363A CELSR2. See Figure S9 for additional data on mutant expression and autoproteolysis. Data are from 4 independent biological replicates. **K,** Same as I, except for full-length WT, L2514A, and M2515A CELSR3. See Figure S9 for additional data on mutant expression and autoproteolysis. Note that the F2511A CELSR3 mutant significantly diminished surface receptor levels, and therefore was excluded from BRET2 analysis. Data are from 5 independent biological replicates. Numerical data are means ± SEM from 3-6 independent biological replicates (depicted as open circles in bar graphs), as indicated in the Figure Legends. Statistical significance was assessed by two-way ANOVA (A) or one-way ANOVA (*, p<0.05; ***, p<0.001). Asterisks depict ANOVA results. See Figure S8 for additional data regarding *Mm* CELSR2 G protein/β-arrestin coupling and Figure S9 for surface expression and autoproteolysis assays for conditions in Figure 7H-K.

To examine this further, we generated a PAR-CELSR2 construct to allow for acute thrombin-mediated TA exposure (Figure 7B and C). We designed PAR-CELSR2 in a similar manner to PAR-LPHN3 by fusing PAR to the CELSR2 CTF so that thrombin-mediated cleavage of PAR at LDPR/S (/denotes cleavage site) would produce the following TA peptide; namely N-SFAVLM…-C (Figure 7B). We subsequently examined the G protein coupling abilities of PAR-CELSR2 following 10-minute 10 U/mL thrombin treatment as before, using PAR-LPHN3*tv7* coupling to Gα13 as a control (Figure 7C). Surprisingly, acute CELSR2 TA exposure alone failed to induce G protein coupling including for GαsS or other G proteins (Figure 7C). While Gα12 exhibited modest changes following thrombin treatment, control experiments revealed that thrombin treatment alone in cells transfected with empty vector can induce the Gα12 TRUPATH BRET2 sensor (Figure S8A). As before, PAR-LPHN3*tv7* displayed strong thrombin-induced Gα13 coupling in the same experiments. Since acute thrombin-mediated TA exposure in PAR-CELSR2 failed to induce G protein coupling in our BRET2 assay, we next tested for Arrestin2/3 (Arr-2/3) induction following acute CELSR2 TA exposure using NanoBiT Arr-2/3 membrane recruitment assays (Figure S8B-D)^60^. Acute CELSR2 TA exposure with the PAR approach also failed to induce Arr-2/3 recruitment (Figure S8C and D), supporting lack of GPCR activation. Altogether, these results suggest that while autoproteolysis is critical for effective *Mm* CELSR2 G protein coupling, acute TA peptide exposure alone using the PAR approach is insufficient for G protein and/or Arr-2/3 induction.

Importantly, the PAR-aGPCR approach generates a “scarred” TA harboring a serine at the N-terminus that is not present in the native TA. While this does not interfere with the LPHN3 TA function, it may impact the CELSR2 TA. Therefore, we utilized another acute TA exposure approach recently developed that generates a native exposed TA^54^. This approach is based on the entirely P’-substrate selectivity of enterokinase, which cleaves following DDDK/^54^. This system has been used to generate functional native exposed TAs for ADGRG6^54^ and LPHN3^61^. We adopted this approach for CELSR2 by capping the native CELSR2 TA (N-TSFAVLM…-C) with SNAP tag-linker-FLAG (Figure 7D). Enterokinase-mediated cleavage in FLAG following DDDK/ exposes the native CELSR2 TA. As a control, we also used a SNAP tag-linker-FLAG-LPHN3*tv*7 in parallel. Surface SNAP tag labelling demonstrated that these fusions were effectively expressed on the cell surface and that enterokinase-cleavage diminished this surface SNAP signal, relative to a C-terminal HA tag (Figure 7E and S8E). We subsequently used this system for BRET2 G protein coupling experiments, including EK-treated cells transfected with empty vector (EV) and 10 nM ISO-treated cells expressing β2-AR as controls (Figure 7F and G). As in Perry-Hauser *et al*., 2022, this approach effectively induced Gα13 coupling with LPHN3 (Figure 7F). However, consistent with our PAR-CELSR2 experiments (Figure 7B and C), acute CELSR2 native TA exposure with this approach failed to induce GαsS coupling alone (Figure 7G).

These results support that CELSRs may display distinct G-protein induction mechanisms compared to LPHN3. To investigate this further, we generated nine point mutations in residues within the CELSR1-3 TA peptide region (Figure 7H). In LPHNs, these residues have shown to be important for autoproteolysis and/or TA-mediated agonism^8, 61^. As before, our CELSR1-3 mutations were generated in full-length *Mm* CELSR1-3 displaying N-terminal HA and C-terminal FLAG tags. We first assessed mutant receptor surface localization and autoproteolysis in HEK293T cells compared to WT forms (Figure S9). CELSR2 F2359A, L2362A, and M2363A mutations impaired autoproteolysis efficiency compared to WT CELSR2 (Figure S9A-D). CELSR1 C2466A, L2469A, and M2470A displayed surface levels and an autoproteolysis profile comparable to WT CELSR1 (Figure S9E, F, I, and J). The CELSR3 F2511A mutation significantly diminished surface expression and localization, while CELSR3 L2514A and M2515A were efficiently expressed on the cell surface and exhibited autoproteolysis comparable to WT CELSR3 (Figure S9G, H, I, and J). We subsequently conducted TRUPATH BRET2 GαsS coupling assays with the CELSR mutants that were effectively expressed on the cell surface (Figure 7I-K). Interestingly, CELSR1 C2466A, L2469A, and M2470A all induced GαsS at a similar intensity to WT CELSR1 (Figure 7I). The three CELSR2 point mutations either abolished (F2359A) or significantly attenuated (L2362A and M2363A) GαsS induction (Figure 7J). However, interpretation of these results in confounded by the observation that these point mutations also impact CELSR2 autoproteolysis. CELSR3 L2514A induced GαsS coupling, while CELSR3 M2515A exhibited a slight reduction in GαsS induction intensity (Figure 7K). Collectively, these results support that these residues are not essential for CELSR1 or CELSR3 GαsS coupling, while CELSR2 requires autoproteolysis in the context of the full-length receptor for robust GαsS induction.

## Discussion

Despite the crucial roles of CELSRs in neurodevelopment, how they function as GPCRs remains unknown. Here, we examined aGPCR:G protein induction principles by focusing on mammalian CELSR1-3 and LPHN3, which represent members of evolutionarily conserved aGPCR families shared across invertebrates to vertebrates. We first examined full-length *Mm* CELSR1-3 autoproteolysis in a heterologous system, and found that full-length *Mm* CELSR1 and CELSR3 are cleavage-deficient, while full-length CELSR2 is cleaved with residue T2357 crucial for efficient autoproteolysis (Figure 1 and 2). We then used an approach to interrogate acute TA exposure-dependent and-independent aGPCR:G protein coupling, first validating this system using *Mm* LPHN3 as a model (Figure 3). Next, we assessed Lphn3 expression together with one C-terminal tail variant in the developing hippocampus, and determined they were highly abundant during postnatal synaptogenesis (Figure 4). Applying the aGPCR:G protein coupling system to *Mm* CELSR1-3, we found that overexpression of full-length *Mm* CELSR1-3 was sufficient to induce G protein coupling, particularly with GαsS (Figure 5). We then found that *Mm Celsr1-3* exhibited distinct, non-overlapping spatial expression profiles during neural circuit assembly (Figure 6). Mechanistically, CELSR2 autoproteolysis was important for effective GαsS coupling, yet acute TA exposure alone was insufficient (Figure 7). Also, mutations in the CELSR1 and CELSR3 TA peptide region failed to block GαsS coupling (Figure 7H-K). Collectively, these results reveal new insights into how CELSRs function as aGPCRs and how aGPCRs mediate G protein coupling induction.

Our study has several inherent limitations which require future experiments to address. First, while even high immunoblot exposures support that the majority of full-length CELSR1 and CELSR3 are uncleaved in our experiments, consistent with their atypical autoproteolysis region (Figure S1D), we cannot exclude the possibility that a small amount of cleavage product is present. Moreover, we were unable to assess native CELSR1-3 autoproteolysis states *in vivo* due to the lack of reliable antibodies against endogenous CELSRs. Future studies will be necessary to circumvent these technical limitations. We employed a panel of 14 G protein BRET2 sensors to assess G protein coupling, but we did not examine CELSR downstream signal transduction pathways. Future studies using a range of biochemical, molecular, and cellular approaches are required to define the precise downstream signaling cascades engaged by CELSR1-3 *in vivo*. Moreover, while our studies focused on mammalian CELSRs, the autoproteolysis and G protein signaling mechanisms of the ancestral invertebrate ortholog (cFMI/Flamingo/starry night) remain to be identified. Interestingly, multiple sequence alignments suggest that cFMI-1 may be cleavage-deficient (Figure 1B) thereby being more comparable to *Mm* CELSR3, but further experiments are required to address this. The PAR/thrombin-mediated acute TA exposure approach also has certain limitations. The endogenous aGPCR TA peptide is protected within the GAIN domain and surrounded by numerous hydrogen bonds and hydrophobic interactions. The PAR-aGPCR fusions remove this molecular context and may alter the native TA conformation. However, our results show robust, thrombin-inducible Gα13 coupling to PAR-Lphn3 as in previous studies^39^, further supporting the validity and effectiveness of this approach for acute TA exposure. Development of approaches with enhanced specificity and temporal resolution for inducible TA peptide exposure, such as the enterokinase (EK) cleavage-mediated strategy, will be of benefit to studies of aGPCR signal transduction.

Our results also prompt multiple new questions. We show in a heterologous system that *Mm* CELSR1/3 exhibit a cleavage-deficient GAIN, while CELSR2 is efficiently cleaved. However, full-length CELSR1-3 all engage GαsS, and effective CELSR2 coupling to GαsS requires autoproteolysis. Interestingly, while autoproteolysis is important for CELSR2:GαsS coupling, acute TA exposure alone using the PAR-CELSR2 or EK-CELSR2 approach is insufficient. This contrasts with LPHN3, where acute TA exposure effectively induces Gα13 coupling in the same experiments using the same approaches (Figure 7). This suggests that autoproteolysis is required in the context of the native protein for CELSR2. One possibility is that the extracellular region plays a crucial role in G protein coupling induction, and therefore acute TA exposure alone is insufficient. Indeed, previous models have been proposed where the extracellular region and 7-transmembrane region communicate, and conformational changes in the extracellular region may directly induce activation without TA exposure^22, 23^. Autoproteolysis may be necessary for the conformational changes allowing communication of the CELSR2 extracellular region with the GPCR.

How do cleavage-deficient aGPCRs, such as CELSR1 and CELSR3, induce G protein coupling in the absence of robust autoproteolysis? A model involving the extracellular region may also help explain this. For CELSR1 and CELSR3, the membrane-proximal “TA” sequence may be still capable of communicating to the 7-transmembrane GPCR via the extracellular region, as shown for other aGPCRs^17, 18^. Thus, certain aGPCRs (mouse/human CELSR1 and CELSR3) have acquired mutations over evolution in the GAIN region rendering them cleavage-deficient yet still capable of inducing G protein coupling, thereby not interfering with their biological functions. However, a typical TA-dependent induction mechanism is unlikely for *Mm* CELSR1 and CELSR3 given that mutations in residues putatively important for TA-mediated agonism fail to block GαsS induction (Figure 7H-K). Unfortunately, we were unable to unequivocally determine the contribution of these residues in CELSR2 towards autoproteolysis-dependent vs. TA-dependent GαsS induction given that point mutations at these sites reduced CELSR2 autoproteolysis (Figure 7J and Figure S9A-D). Nonetheless, our studies support that cleavage-independent mechanisms and several modes of signal transduction exist for aGPCRs.

What are the biological roles of CELSR1-3 in the mammalian brain? Critical insights into *C. elegans* and *Drosophila* Flamingo/starry night function have been determined in invertebrate systems, and Flamingo has been associated with the planar cell polarity pathway in development^40–52^. These previous invertebrate studies support that CELSRs play integral roles in circuit wiring and brain development. Intriguingly, important studies have shown that CELSR2 and CELSR3 modulate opposing roles in mammalian neurite development in cultured neurons^63^. Our studies support differential spatial expression patterns of *Celsr1-3* during neural circuit assembly (Figure 6). These results show that *Celsr2/3* are the predominant isoforms and are expressed in neurons through postnatal life, supporting a neuronal role of *Celsrs* during mammalian postnatal development. *Celsr2/3* also partially overlap in the hippocampus, and therefore we speculate that a cleaved (*Celsr2*) and cleavage-deficient (*Celsr3*) isoform may have functions in the same cells. Cleavage-deficient *Celsr1* was restricted to early development, and persisted in the region responsible for granule cell adult neurogenesis, suggesting a distinct biological role from *Celsr2/3*. Therefore, *Celsr1-3* likely have non-overlapping functions. Future *in vivo* functional studies are required to address this. Our studies elucidating CELSR1-3 autoproteolysis and G protein coupling mechanisms provide a framework for future studies of CELSR function *in vivo*.

In summary, we investigated autoproteolytic cleavage and Gαs coupling mechanisms of vertebrate *Mm* CELSR1-3. We demonstrated that *Mm* CELSR1-3 display different autoproteolytic cleavage profiles and distinct autoproteolysis-dependent G protein induction paradigms. Thus, aGPCRs display several modes of G protein engagement. Overall, these studies expand our current understanding of aGPCR molecular mechanisms and open new avenues of future investigation into CELSR biological function.

### Limitations of the Study

Here, we utilized a heterologous system to examine full-length mouse CELSR1-3 autoproteolysis upon overexpression in HEK293T cells. We were unable to investigate native mouse CELSR1-3 autoproteolysis in brain tissue due to the lack of reliable antibodies together with lack of CELSR1-3 KO brain homogenate to serve as a control. Future studies are required to determine native CELSR1-3 autoproteolysis *in vivo*.

## Methods

### KEY RESOURCES TABLE

| REAGENT or RESOURCE | SOURCE | IDENTIFIER |
| --- | --- | --- |
| **Antibodies** |  |  |
| Anti-HA mouse | Covance/Biolegend | MMS101R |
| Anti-HA rabbit | Cell Signaling Technologies | 3724 |
| Anti-NeuN mouse | EMD Millipore | MAB377 |
| Goat anti-Mouse Alexa Fluor 488 | ThermoFisher | A11001 |
| Goat anti-Mouse Alexa Fluor 546 | ThermoFisher | A11003 |
| Goat anti-Mouse Alexa Fluor 647 | ThermoFisher | A21236 |
| Goat anti-rabbit Alexa Fluor 488 | ThermoFisher | A11034 |
| Goat anti-rabbit Alexa Fluor 546 | ThermoFisher | A11010 |
| Goat anti-rabbit Alexa Fluor 647 | ThermoFisher | A21245 |
| Goat anti-chicken Alexa Fluor 647 | ThermoFisher | A21449 |
| Goat anti-guinea pig Alexa Fluor 647 | ThermoFisher | A21450 |
| HRP-conjugated goat anti-mouse secondary antibody | Biotium | 20400-1mL |
| Phalloidin Alexa 647 | ThermoFisher | A22287 |
| Phalloidin-iFluor 405 | Abcam | Ab178752 |
| Anti-FLAG iFluor488 conjugate | Genscript | A01809 |
| Anti-HA iFluor647 conjugate | Genscript | A01808 |
| Opal 520 | Akoya Biosciences | OP-001001 |
| Opal 570 | Akoya Biosciences | OP-001003 |
| Opal 690 | Akoya Biosciences | OP-001006 |
| **Bacterial and Virus Strains** |  |  |
| DH10β | ThermoFisher | 18297010 |
| Stellar cells | Takara |  |
| **Chemicals, Peptides, and Recombinant Proteins** |  |  |
| Bovine Serum Albumin Fraction V | Roche | 10735086001 |
| Normal Goat Serum | Jackson ImmunoResearch | 005000121 |
| Bovine Serum Albumin | Sigma | A3803 |
| Coelenterazine-400a | NanoLight Technologies | 340 |
| DAPI | Roche | 10236276001 |
| DMEM | Gibco | 11995065 |
| Fetal Bovine Serum | Gibco | 16000044 |
| Hanks' balance Salt Solution | Gibco | 14175095 |
| HEPES | Sigma | H3375 |
| Matrigel Membrane Matrix | ThermoFisher | CB-40234 |
| Poly-D-lysine | Gibco | A38904-01 |
| MEM Non-essential Amino Acid Solution | Sigma | M7145 |
| Nano-Glo Live Cell Assay System | Promega | N2011 |
| Opti-MEM | Gibco | 31985070 |
| Paraformaldehyde | Electron Microscopy Science | 15714 |
| Penicillin/Streptomycin | Corning | MT30002CI |
| Thrombin | Sigma | T4846 |
| Isoproterenol | Sigma | I6504 |
| TransIT-2020 | Mirus | MIR5400 |
| LipoD293 | SignaGen Laboratories | SL100668 |
| Versene | Gibco | 15040066 |
| Enterokinase, light chain | NEB | P8070S |
| SNAP-Surface Alexa Fluor 488 | NEB | S9129S |
| Mouse *Adgrl3* RNAscope probe for Accession #NM_001359828.1 region 4660-5680 (Ct-Lphn3) | Advanced Cell Diagnostics | Mm-Adgrl3-O2-C2<br>#1046031-C2 |
| Mouse *Adgrl3* RNAscope probe for Accession #NM_198702.2 region 2338-3312 | Advanced Cell Diagnostics | Mm-Lphn3-C3<br>#317481-C3 |
| Mouse *Celsr1* RNAscope probe for Accession #NM_009886.2 region 4742-5601 | Advanced Cell Diagnostics | Mm-Celsr1<br>#317931 |
| Mouse *Celsr2* RNAscope probe for Accession #NM_001004177.2 region 3910-4817 | Advanced Cell Diagnostics | Mm-Celsr2-C2<br>#317941-C2 |
| Mouse *Celsr3* RNAscope probe for Accession #NM_080437.2 region 7034-7885 | Advanced Cell Diagnostics | Mm-Celsr3-C3<br>#319241-C3 |
| RNAscope Multiplex Fluorescent manual assay | Advanced Cell Diagnostics | 323100 |
| RNA-protein Co-detection ancillary kit | Advanced Cell Diagnostics | 323180 |
| To-Pro | ThermoFisher | T3605 |
| 0.45 µm pore nitrocellulose membrane | Bio-Rad | 1620251 |
| Kaleidoscope protein standard | Bio-Rad | 1610375 |
| PageRuler protein standard | Thermo | 26619 |
| Prolong Gold Antifade Reagent | Invitrogen | P36930 |
| **Experimental Models: Cell Lines** |  |  |
| HEK293T | ATCC | CRL-11268 |
| GKO HEK293 | Kind gift from Asuka Inoue, Tokyo University | Alvarez-Curto *et al.*, 2016, J Biol Chem. ; Grundmann *et al.*, 2018, Nat Commun. |
| Arr-2/3 KO HEK293 | Kind gift from V.V. Gurevich, Vanderbilt University | Alvarez-Curto *et al.*, 2016, J Biol Chem. Grundmann *et al.*, 2018, Nat Commun. |
| **Experimental Models: Organisms/Strains** |  |  |
| C57/BL6J | Jackson Laboratories | 000664 |
| **Oligonucleotides** |  |  |
| Various cloning primers | IDT | N/A |
| **Recombinant DNA** |  |  |
| pEB PAR-Lphn3tv5 | This study |  |
| pEB PAR-Lphn3tv7 | This study |  |
| pEB PAR-Lphn3tv9 | This study |  |
| pEB HA-Celsr1-FLAG | This study |  |
| pEB HA-Celsr1 A2464T-FLAG | This study |  |
| pEB HA-Celsr2-FLAG | This study |  |
| pEB HA-Celsr2 T2357A-FLAG | This study |  |
| pEB HA-Celsr2 T2357G-FLAG | This study |  |
| pEB HA-Celsr3-FLAG | This study |  |
| pEB PAR-Celsr2 | This study |  |
| pEB SNAP-FLAG-Lphn3<sup>tv7</sup>-HA | This study |  |
| pEB SNAP-FLAG-Celsr2-HA | This study |  |
| pEB HA-Celsr1 C2466A-FLAG | This study |  |
| pEB HA-Celsr1 L2469A-FLAG | This study |  |
| pEB HA-Celsr1 M2470A-FLAG | This study |  |
| pEB HA-Celsr2 F2359A-FLAG | This study |  |
| pEB HA-Celsr2 L2362A-FLAG | This study |  |
| pEB HA-Celsr2 M2363A-FLAG | This study |  |
| pEB HA-Celsr3 F2511A-FLAG | This study |  |
| pEB HA-Celsr3 L2514A-FLAG | This study |  |
| pEB HA-Celsr3 M2515A-FLAG | This study |  |
| TRUPATH kit | Addgene | 1000000163 |
| pcDNA3.1 FLAG-β<sub>2</sub>-Adrenergic receptor | Addgene | 14697 |
| pCMV-hsLphn3 | Lu *et al.*, 2015, Structure |  |
| Membrane-anchored LgBiT (AG10-CAAX) | Spillman *et al.*, 2020 |  |
| pEB *Mm*-Arr2-SmBiT | This study |  |
| pEB *Mm*-Arr3-SmBiT | This study |  |
| pEB *Mm* GRK2 | This study |  |
| **Software** |  |  |
| SnapGene | GSL Biotech |  |
| NIS-Elements AR | Nikon |  |
| ImageJ | National Institutes of Health |  |
| Adobe Photoshop | Adobe |  |
| Adobe Illustrator | Adobe |  |
| Graphpad Prism 8.0, 9.0 | Graphpad |  |

### RESOURCE AVAILABILITY

#### Lead contact

Further information and requests for resources and reagents should be directed to and will be fulfilled by the lead contact, Richard C. Sando.

#### Materials availability

All materials generated in this study will be openly shared upon request free of charge.

#### Data and code availability

Original western blot images have been submitted with the manuscript. No original code has been generated in this study.

### METHODS DETAILS

#### Mice

Mice were weaned at 18-21 days of age and housed in groups of 2 to 5 on a 12 h light/dark cycle with food and water *ad libidum*. Vanderbilt Animal Housing Facility: All procedures conformed to National Institutes of Health Guidelines for the Care and Use of Laboratory Mice and were approved by the Vanderbilt University Administrative Panel on Laboratory Animal Care. Mice for RNA *in situ* studies were euthanized at indicated developmental time-points (postnatal day 5, 10, or 21) and whole brains were flash frozen by indirect exposure to liquid N_2_, followed by the procedure for fresh frozen tissue outlined below.

#### Cell Lines

GKO HEK293 cells used for all BRET2 assays were originally a kind gift from Asuka Inoue (Tokyo University, Japan) (Alvarez-Curto *et al*., 2016; Grundmann *et al*., 2018); and were provided to our studies as a kind gift from Drs. Vsevolod Gurevich and Chen Zheng. GKO HEK293 cells were maintained in DMEM (Gibco Cat# 11995065) containing 10% FBS (Gibco Cat# 16000044), 1X Penicillin-Streptomycin (Corning Cat# MT30002Cl), and 1x MEM Non-essential Amino Acid (NeA) Solution (Sigma Cat# M7145) at 37°C and 5% CO_2_. HEK293 Arrestin2/3 KO used for Indirect Arrestin assays were originally generated by Milligan lab from University of Glasgow (Alvarez-Curto *et al*., 2016) and were a kind gift from Drs. Vsevolod Gurevich and Chen Zheng (Vanderbilt University). HEK293T cells (ATCC #CRL-11268) were used to examine cDNA overexpression and protein cell surface localization. Cells were maintained in DMEM (Gibco Cat# 11995065) plus 10% FBS (Gibco Cat# 16000044) and 1X Penicillin-Streptomycin (Corning Cat# MT30002Cl) at 37°C and 5% CO_2_ for a maximum of 20 passage numbers.

#### Plasmids

All full-length *Mm* CELSR1-3, PAR fusion, and SNAP-FLAG fusion overexpression cDNAs were encoded in the pEB Multi-Neo vector (Wako Chemicals, Japan). All PAR constructs contained an N-terminal fusion of the human thrombin receptor PAR1 signal peptide and PAR cleavage sequence N-MGPRRLLLVAACFSLCGPLLSARTRARRPESKATNATLDPRS-C. PAR-LPHN3 and PAR-CELSR constructs were designed according to Mathiasen *et al*., 2020. For PAR-LPHN3, the PAR sequence was fused immediately upstream of the tethered agonist sequence beginning at N-FAVLMAH…-C. Thrombin-mediated cleavage of the PAR at N-…LDPR/S-C (/ denotes cleavage site) exposes the following tethered agonist sequence (N-SFAVLMAH…-C). *Mus musculus* PAR-LPHN3 variants Lphn3*tv*5 (NCBI Reference Sequence NM_001347371.2), Lphn3*tv*7 (NCBI Reference Sequence NM_001359828.1), and Lphn3*tv*9 (NCBI Reference Sequence NM_001359830.1) were cloned into pEB Multi-Neo via gBlocks (Integrated DNA technologies). *Mus musculus* PAR-CELSR2 constructs were based CELSR2 Uniprot #Q9R0M0 and synthesized via codon-optimized gBlock fragments (PAR-CELSR2 amino acids #2359-2919 of *Mm* CELSR2). Full-length *Mm* CELSR1 and *Mm* CELSR2 were cloned from postnatal day 21 mouse brain cDNA library produced in-house using Superscript IV Reverse Transcriptase (Invitrogen #18091050). *Mm* Celsr1 corresponded to NCBI Reference Sequence NM_009886.2 and Uniprot #O35161. *Mm* Celsr2 corresponded to NCBI Reference Sequence NM_017392.4 and Uniprot #Q9R0M0. *Mm* Celsr3 was codon-optimized and synthesized at Gene Universal Inc. (Newark, DE) and corresponded to Genbank #AIJ27736.1. The native signal peptide based on Uniprot was replaced with a preprotrypsin leader and HA tag (N-MSALLILALVGAAVAYPYDVPDYA-C). cDNAs contained the N-terminal preprotrypsin leader-HA and C-terminal FLAG tags. The ADGRL3 construct used as a positive control in cleavage assays consisted of full-length human LPHN3 inserted into pCMV5 using restriction cloning with an N-terminal FLAG tag and an HA tag introduced into the first extracellular loop of the seven-transmembrane region (Lu *et al*., 2015). The SNAP-FLAG-Lphn3*tv*7 and SNAP-FLAG-CELSR2 constructs contained the following sequence of features fused upstream of the native tethered agonist sequence: IgK signal peptide (METDTLLLWVLLLWVPGSTGDAGAQ) – SNAP-tag – linker (GGSGGSGGS) – FLAG (DYKDDDDK). Enterokinase-mediated cleavage following DDDDK/ was designed to expose the native TA sequence for Lphn3*tv*7 (TNFAVLMAH…) and CELSR2 (TSFAVLMDM…). Both constructs were cloned into pEB and contained a C-terminal HA-tag fusion. The TRUPATH kit was purchased from Addgene (#1000000163) and was a gift from Bryan Roth. The pcDNA3 FLAG *H. sapiens* beta-2-adrenergic receptor used as a control in BRET2 experiments was a gift from Robert Lefkowitz (Addgene plasmid #14697). *Mm* Arr-2 (NP_796205.1) and *Mm* Arr-3 (NP_001258289.1) were cloned via gBlocks (Integrated DNA technologies) to create N-terminal fusions of SmBiT in pEB-Multi-Neo. *Mm* GRK2 (NP_001277747.1) was cloned from gBlocks (Integrated DNA Technologies) into pEB-Multi-Neo. Cells transfected with empty pEB-Multi-Neo were used as negative control conditions in all experiments. All molecular cloning was conducted with the In-Fusion Assembly system (Takara #638948).

#### Antibodies

The following antibodies and reagents were used at the indicated concentrations for immunocytochemistry: anti-HA mouse (Covance Cat# MMS101R; 1:1,000), anti-HA rabbit (Cell Signaling Technologies Cat# 3724; 1:2,000), Alexa Fluor 647 Phalloidin (Invitrogen Cat# A22287; 1:40 diluted in methanol), and corresponding fluorescently-conjugated goat secondary antibodies from Life Technologies (1:1,000). For autoproteolysis assays, anti-FLAG antibody conjugated to iFluor 488 (GenScript A01809) at 1:2,000 dilution, or anti-HA antibody conjugated to iFluor 647 (GenScript A01808) at 1:1,000 were used. For double IHC/RNA in situ experiments, anti-NeuN mouse (EMD Millipore #MAB377) was used at a 1:500 dilution and detected with HRP-conjugated goat anti-mouse secondary antibody (Biotium #20400-1mL) at 1:500 together with Opal 520 reagent at 1:1,000 (Akoya Biosciences #OP-001001).

#### BRET2 assays

HEK G protein K.O cells (Grundmann *et al*., *Nature Commun.*, 2018) were plated into 12-well plate at a density of 3-4 × 10^5^ cells in 1 mL per well. HEK G K.O. media contained 1x DMEM (Gibco Cat# 11995065) plus 10% FBS (Gibco Cat#16000044), 1X Penicillin-Streptomycin (Corning Cat#MT30002Cl) with 1x MEM Non-essential Amino Acid (NeA) Solution (Sigma Cat# M7145). After 16-24 hours, cells were co-transfected with receptor of interest and TRUPATH plasmids at 1:1:1:1 DNA ratio (receptor:Gα-RLuc8:Gβ:Gγ-GFP2) via TransIT-2020 (Mirus Cat# MIR5400). Each condition required 97 µL of room-temperature 1x Opti-MEM (Gibco Cat# 31985070), 1 µL each DNA plasmid at 1µg/µL concentration), and 3 µL of room-temperature and gently-vortexed TransIT-2020 reagent. The TransIT-2020:DNA complexes mixture were gently mixed via pipetting 10 times and incubated at room temperature for 20 mins before adding drop-wise in the well. The plate was rocked gently side to side and incubated at 37°C 24 hours before harvesting. In each well, media was aspirated, and cells were washed with 1 mL warm PBS. Cells were detached with 300 µL warm Versene (Gibco Cat# 15040066) and incubated at 37°C for 5 mins then resuspended via pipetting 10 times. Cells were plated in complete DMEM containing 1x NeA at 200 µL with a density of 30,000–50,000 cells per well in Matrigel-coated 96-well assay plate. Each experimental condition was plated into three separate wells within the 96-well assay plate. BRET2 assays were performed 48 hours after transfection. In each well, media was aspirated and cells were incubated in 80 µL of 1x Hanks’ balanced Salt Solution (Gibco Cat# 14175095) with 20 mM HEPES (Sigma Cat# H3375, pH 7.4) and 10 µL 100 µM Coelenterazine-400a (NanoLight Technologies Cat# 340) diluted in PBS for 5 minutes before adding 10 µL of vehicle solution, isoproterenol agonist (Sigma #I6504), or Thrombin (Sigma Cat# T4648) at 100 U/mL concentration for a final concentration of 10 U/mL, or 10 µL of Thrombin vehicle, which is 0.1% BSA (Roche Cat# 10735086001) in PBS. After 10 mins of incubation, BRET intensities were measured via BERTHOLD TriStar^2^ LB 942 Multimode Reader with Deep Blue C filter (410nm) and GFP2 filter (515 nm). The BRET ratio was obtained by calculating the ratio of GFP2 signal to Deep Blue C signal per well. The BRET2 ratio of the three wells per condition were then averaged. Net BRET2 was subsequently calculated by subtracting the BRET2 ratio of cells expressing donor only (Gα-RLuc8) from the BRET2 ratio of each respective experimental condition. Net BRET2 differences were then compared as described in the Figures, by subtracting Net BRET2 ratios of induced (thrombin/isoproterenol) conditions from vehicle treated conditions, or conditions overexpressing indicated receptors compared to empty vector (EV). For plasmid copy-dependent BRET2 experiments, conditions were transfected in a 12-well plate format with the same total amount of plasmid DNA and varying copies of experimental plasmid, adjusted to the same total amount with empty vector (pEB-multi). For Enterokinase treatment BRET2 assays, GKO HEK293 cells were plated and transfected as above. On the day of the assay, media in each well was aspirated and cells were incubated in 80 μL of 1x Hanks’ balanced Salt Solution with 20 mM HEPES and 10 μL 100 μM Coelenterazine-400a diluted in PBS for 5 minutes before adding either 10 μL of 5.5 U of Enterokinase (New England Biolabs #P8070S) in PBS or 10 μL of vehicle PBS. After 15 mins incubation, BRET2 intensities were measured as above.

#### Adhesion GPCR autoproteolysis assays

The autoproteolysis assay was performed similarly to previously described (Araç *et al*., 2012). HEK293T cells were cultured in Dulbecco’s Modified Eagle Medium (Gibco 11965118) + 10 % fetal bovine serum (Sigma F0926) at 37 °C in a 5 % CO_2_ atmosphere. At 70 % confluency, cells were transfected with 2 micrograms of either CELSR constructs or empty vector DNA in a 6-well plate (Fisher Scientific FB012927) using a 1:3 mixture of LipoD293 (SignaGen Laboratories SL100668). 48 hours later, the cells were washed with 1 mL of Dulbecco’s Phosphate Buffered Saline (PBS) (Gibco 14190144) and the PBS was aspirated; dry adhered cells were placed at-80 °C. Cells were thawed the following day and the following steps were performed at 4 °C: cells were resuspended in a solution of PBS + 0.01 % bovine serum albumin (BSA) (Sigma A3803) and centrifuged using a swinging-bucket rotor at 2,600 x g for 5 minutes. The supernatant was aspirated and the cell pellet was resuspended in 500 µL of solubilization buffer (20 mM HEPES pH 7.4, 150 mM NaCl, 2 mM MgCl_2_ 0.1mM EDTA, 2 mM CaCl_2_, 1% (v/v) Triton X-100). The resuspended cell pellet was incubated at 4 °C with rotation for 30 minutes for solubilization. The solubilized cell pellet was then centrifuged at 20,000 x g for 15 minutes and the supernatant was collected as the solubilized cell lysate fraction. Cell lysates were run on 12 % SDS-PAGE gels and transferred to 0.45 µm pore nitrocellulose membranes (Bio-Rad 1620251) using a wet transfer system (Bio-Rad) for 1 hour at 100 V in a transfer buffer of 24.9 mM Tris-HCl and 193 mM glycine with 20 % (v/v) methanol. Either the Kaleidoscope (Bio-Rad 1610375) or the PageRuler Plus (Thermo 26619) protein standards were used. After transfer, the nitrocellulose membrane was blocked for one hour at room temperature using 4 % BSA in TBST (20 mM Tris Base pH 7.4, 150 mM NaCl, 0.1 % (v/v) Tween-20). After blocking, membranes were incubated with either anti-FLAG antibody conjugated to iFluor 488 (GenScript A01809) at 1:2,000 dilution, or anti-HA antibody conjugated to iFluor 647 (GenScript A01808) at 1:1,000 dilution overnight at 4 °C with gentle rocking. The next day, membranes were washed 4 times with TBST for 5 minutes each at room temperature. The membranes were imaged at respective wavelengths using the ChemiDoc imaging system (Bio-Rad).

#### RNA in situ hybridizations

**Tissue Preparation and Sectioning:** Wild-type C57BL/6J (Jackson #000664) mice were taken from their home cages at postnatal day (P) 5, P10, and P21, and whole brain tissue was collected in the following manner. Brains were rapidly dissected following brief anesthesia with either ice (P5) or isoflurane (P10 and P21) and placed in a rectangular cryomold (Epredia Peel-a-way #18-30) which was flash-frozen in liquid N_2_ for 15 seconds to allow for indirect exposure of the tissue with liquid N_2_. The brain was subsequently embedded in O.C.T. Compound (Fisher Healthcare, #4585) within a second cryomold using a bath of 2-methylbutane (Sigma-Aldrich, #M32631-46) equilibrated with dry ice. Once frozen, the blocks were stored at-80°C until cryosectioning.

The frozen blocks were removed from the-80°C freezer and allowed to sit in the cryostat (Leica CM 1950, #047742456) at-20°C for 1 hour to equilibrate. The microtome blade for slicing (Sakura, #4689), forceps (Fine Science Tools), razor blade for block trimming, and paintbrushes for manipulating sections, and the antiroll plate (Leica, #14047742497) were also all placed in the cryostat and allowed to equilibrate. Tissue was sectioned at 15 µm and mounted directly onto room temperature Diamond White Glass microscope slides (Globe Scientific Inc., white frosted 25×75×1mm, charged +/+, #1358W). Once mounted, the slides were kept in the cryostat until all sectioning was complete. Sections were subsequently dried at-20°C for 1 hour, then stored at-80°C.

Tissue Pre-Treatment: The RNAscope Multiplex Fluorescent manual assay (Advanced Cell Diagnostics #323100) was carried out using the Fresh Frozen sample preparation according to the manufacturer’s protocol as described below. The RNAscope® Hydrogen Peroxide (Advanced Cell Diagnostics, #322335) and RNAscope® Protease IV (Advanced Cell Diagnostics, #322336) reagents were set out on the benchtop to equilibrate to room temperature. Sections were removed from the-80°C freezer and placed immediately into ice cold 4% PFA (Electron Microscopy Sciences, #15714)/PBS (MP Biomedicals, 1 tab/100 mL, #2810306) within a glass slide holder (Epredia RA Lamb Glass Coplin Jar, Fisher #E94). Slides were incubated at 4°C for 15 minutes to fix the tissue and subsequently washed twice with 1x PBS. Slides were subsequently dehydrated in the following ethanol (Decon Laboratories, Inc., 200 proof, #2705HC) gradient: 50% EtOH/ddH_2_O for 5 minutes, 70% EtOH/ddH_2_O for 5 minutes, followed by two treatments with 100% EtOH (50 mL of each treatment). After the final 100% step, the slides were placed section side up on a paper towel and allowed to air dry for 5 minutes. Then a hydrophobic pen (IHC World, super pap pen, #SPR0905) was used to draw a barrier around each section, which dried at room temperature for 5 minutes. While the barriers were drying, the HybEZ™ Humidity Control Tray with lid (Advanced Cell Diagnostics, #310012) was prepared. A sheet of HybEZ™ Humidifying Paper (Advanced Cell Diagnostics, #310025) was placed on the bottom of the tray and sprayed with ddH_2_O until damp. The EZ-Batch™ Slide Holder (Advanced Cell Diagnostics, #310017) was placed inside the humidity control tray and the slides were placed in the holder. Three drops of hydrogen peroxide were added to each section. The cover was placed over the humidity control tray and the slides were left to incubate for 10 minutes at room temperature. Once the incubation was complete, the slides were washed twice with ddH_2_O, removing excess liquid after each wash via a vacuum. Slides were reinserted into the slide holder and 4 drops of Protease IV were added to each section, followed by incubation at room temperature for 30 minutes. While the slides were incubating the RNAscope® Probes were prepared in the following manner. First the C2 (Advanced Cell Diagnostics: Mm-Adgrl3-O2-C2 Mus musculus Adhesion G protein-coupled receptor L3 (Adgrl3) transcript variant 7 mRNA, #1046031-C2) and C3 (Advanced Cell Diagnostics, Mm-Adgrl3-C3, #317481-C3) probes and 100 µl/sample of the RNAscope® Probe Diluent (Advanced Cell Diagnostics, #300041) were placed in a heat block (Thermo Scientific, #88870003) at 40°C for 10 minutes, and subsequently removed from the heat and incubated at room temperature for 10 minutes. The probes were combined to form a probe mix consisting of 100 µl of probe diluent per section, and 2 µl each of the C2 and C3 probes per section, respectively. The HybEZ™ II Oven (Advanced Cell Diagnostics, #321720) was then pre-warmed to 40°C. Once the Protease IV incubation was complete, the slides were washed twice with 1x PBS, placed back in the slide holder and humidity control tray and 100 µl of probe mix were added to each section. The tray was placed in the HybEZ™ Oven for 2 hours at 40°C to hybridize the probes. While the incubation was occurring, the 1x RNAscope® wash buffer solution (Advanced Cell Diagnostics, #310091) and 5x Saline Sodium Citrate (SSC) buffer was prepared. The 20x SSC stock contained175.3 g of NaCl (Fisher Chemical, certified ACS, crystalline, #S271-1) and 88.2 g of sodium citrate (Fisher Chemical, dihydrate, granular, #S279-500) in ddH_2_O, pH 7.0. Following the 2-hour period slides were washed twice with 1x RNAscope wash buffer and placed in 5x SSC buffer overnight at 4°C.

RNAscope® Multiplex Fluorescent Assay: The following reagents were equilibrated at room temperature for 1 hour: RNAscope® Multiplex FL v2 AMP1 (Advanced Cell Diagnostics, 323101), RNAscope® Multiplex FL v2 AMP2 (Advanced Cell Diagnostics, 323102), RNAscope® Multiplex FL v2 AMP3 (Advanced Cell Diagnostics, 323103), RNAscope® Multiplex FL v2 HRP C1 (Advanced Cell Diagnostics, 323104), RNAscope® Multiplex FL v2 HRP C2 (Advanced Cell Diagnostics, 323105), RNAscope® Multiplex FL v2 HRP C3 (Advanced Cell Diagnostics, 323106) and RNAscope® Multiplex FL v2 HRP Blocker (Advanced Cell Diagnostics, 323107). While this equilibration was occurring, the HybEZ™ Oven was equilibrated to 40°C. Slides were removed from 5x SSC and washed twice with RNAscope® wash buffer. Three drops of the AMP1 were applied to each section and the humidity control tray was placed back in the oven where it incubated for 30 minutes at 40°C. Slides were subsequently washed twice with wash buffer and placed back in the slide holder and 3 drops of AMP2 were applied to each section and left to incubate in the oven at 40°C for 30 minutes. Slides were washed twice, and treated with 3 drops of AMP3 at 40°C for 15 minutes. While the AMP3 incubation was occurring, the dye solutions were prepared as follows. The green and red dyes were prepared at a concentration of 1:1000 by combining 1000 µl of TSA Buffer (Advanced Cell Diagnostics, 322809) with 1 µl of Opal 520 Reagent (in DMSO, Akoya Biosciences, OP-001001) and Opal 570 Reagent (in DMSO, Akoya Biosciences, OP-001003) respectively.

Once the 15-minute AMP3 incubation was complete the slides were washed and inserted back into the slide holder, then 3 drops of HRPC1 were applied to each slide, and that was allowed to incubate for 15 minutes at 40°C. Once that was complete the slides were washed and put back into the slide holder and 3 drops of the HRP Blocker were added to each section. This was allowed to incubate for 15 minutes at 40°C. Slides were washed and 150 µl of the Opal 570 dye mix was added to each section. This was allowed to incubate at 40°C for 30 minutes. This process was then repeated for the C3 channel, which was treated with Opal 520 dye.

Counterstaining Mounting and Imaging: Four drops of RNAscope® DAPI (Advanced Cell Diagnostics, #323108) were added to each section for the purpose of counterstaining and left to sit at room temperature for 30 seconds. The DAPI was gently tapped off the slide and 50 µl of Prolong™ Gold antifade reagent (Invitrogen, #P36930) was added inside the barrier but not directly touching the section, avoiding bubbles. A glass coverslip (Corning, 24 × 60 mm, #2975-246) was lowered onto the slide, slides were allowed to dry overnight in a dark slide box (Fisher Brand, #03-448-4) at 4°C before imaging. Slide boxes were stored in the cold room a 4°C for long term storage. Slides were imaged on a Nikon Eclipse T*i*2 Confocal Microscope within 1 week of RNAscope® completion. Images were taken at 10x and 60x magnification.

Images were analyzed using NIS-Elements AR 5.41.02 software in the following manner. Using the Automated Measurements and Automated Measurements Results functions, thresholds were assigned to the 405, 488, 561, and 640 channels. This was done using the per channel mode and with the following parameters: (for 405) smooth, clean, separate, and fill holes off, and the size option selected with lower bound of 5 µm; (for 488, 561, and 640) smooth, clean, separate, and fill holes off, no size restrictions. Once the threshold had been set for each channel, the data was exported to excel using the software’s built in export feature. The sum intensity for all points for a given channel was found, along with the total pixel area occupied of each channel. Then the sum intensity of each channel was divided by the DAPI area for its respective image to produce a ratio of the sum intensity of the gene-of-interest to the area occupied by internal control DAPI. For each region and postnatal day, 5-10 separate images were analyzed and subsequently averaged to generate an ‘n’ value for each mouse. Three separate mice were analyzed for each postnatal age, and quantitative data depicts the average values from three mice.

#### Double immunohistochemistry/RNA in situ hybridizations

NeuN/RNA *in situ* experiments were conducted in the following manner, essentially as described in the manufacturer’s protocol (Advanced Cell Diagnostics #323180 and #323100). Tissue collection, sectioning, and pretreatment was conducted as described above for standard RNA *in situ* experiments up until the initial 10-minute room temperature hydrogen peroxide treatment. Following hydrogen peroxide treatment, slides were washed twice with ddH_2_O followed by once with 1x-PBS-T (PBS with 0.1% Tween-20). The slides were placed back in the slide holder and 150 µL of the primary antibody (anti-NeuN Mouse, EMD Millipore Corp., #MAB377) diluted in RNAscope® Co-Detection Antibody Diluent (Advanced Cell Diagnostics, #323160) in a 1:500 concentration was added to each section. Slides were incubated at 4°C overnight in the humidity control tray. Post-primary Fixation and Protease Treatment: After incubation with the primary antibody, slides were washed three times with 1x-PBS-T at room temperature. Then slides were submerged in 10% Neutral Buffered Formalin (Sigma-Aldrich, #65346-85) for 30 minutes at room temperature. Following that incubation, slides were washed four times with PBS-T. Slides were subsequently placed back into the humidity control tray, and 4 drops of Protease 4 were added and incubated for exactly 30 minutes at room temperature. After incubation the slides were washed three times with ddH_2_O. The RNAscope Multiplex fluorescent assay was then performed, as described above. Following the last HRP blocker step in the RNAscope Multiplex assay, immunofluorescence for NeuN was performed. HRP-conjugated goat anti-mouse secondary antibody (Biotium #20400-1mL) was diluted in Co-Detection Antibody Diluent (Advanced Cell Diagnostics, #323160) at a 1:500 concentration was added to completely cover the sections and allowed to incubate at room temperature for 30 minutes. Slides were subsequently washed twice with 1x PBS-T. Then 150 µL of the previously prepared Opal dye (Akoya Biosciences) was added to the slides and incubated for 10 minutes at room temperature. Then the slides were washed twice with 1x PBS-T and ready for counterstaining and mounting as described above for standard *in situs*.

#### Surface labeling and immunocytochemistry

Cover glass (#0, 12 mm, Carolina Biological Supply Company #633009) was placed into 24-well plates and coated for 2 hours with 100 µL of 50 µg/mL poly-D-lysine (Gibco #A38904-01) in the 37°C tissue culture incubator. Excess poly-D-lysine was removed, coverslips were washed 3x with sterile ddH_2_O, and dried for 30-minutes. HEK293T cells were plated at 1.5-2 × 10^5^ cells/well in 0.5 mL complete DMEM. After 16-24 hours, cells were transfected with indicated experimental plasmid via TransIT-2020 (Mirus MIR5400) with a total of 0.5 µg DNA amount/condition/well. After 48-hrs post-transfection, cells were washed briefly once with PBS, fixed with 4% PFA (Electron Microscopy Science Cat# 15714)/4% sucrose/PBS for 20 min at 4°C, and washed 3 × 5 minutes in PBS. For surface receptor labeling of HA tag, samples were then transferred directly into blocking buffer (4% BSA (Sigma Cat# 10735086001)/3% goat serum (Jackson Immunoresearch #005000121)/PBS). For total receptor labeling, samples were permeabilized in 0.2% Triton X-100/PBS for 5 minutes at room temperature and then transferred to blocking buffer. Samples were incubated in blocking buffer for 1 hour, and subsequently incubated with diluted primary HA tag antibody (anti-HA rabbit (Cell Signaling Technologies Cat# 3724; 1:2,000)) in blocking buffer for 2 hours at room temperature. Samples were then washed 3 × 5 minutes in PBS, incubated with diluted fluorescently-conjugated secondary antibody (goat anti-rabbit IgG Alexa Fluor 488 Cat#A11032; 1:1,000) together with fluorescently-conjugated Alexa Fluor 647 Phalloidin (Invitrogen Cat# A22287; 1:40 diluted in methanol) and DAPI (Sigma Cat# 10236276001; 1:1,000) in blocking buffer for 30 minutes, washed three times in PBS, and mounted on UltraClear microscope slides (Denville Scientific Cat# M1021) using 10 µL ProLong Gold antifade reagent (Invitrogen, #P36930) per coverslip. Imaging regions-of-interest were chosen at random. ‘Low-magnification’ images were collected with a 20x objective and ‘high-magnification’ images a 60x objective (see Imaging section for details).

#### Imaging

Images were acquired using a Nikon A1r resonant scanning Eclipse Ti2 HD25 confocal microscope with a 10x (Nikon #MRD00105, CFI60 Plan Apochromat Lambda, N.A. 0.45), 20x (Nikon #MRD00205, CFI60 Plan Apochromat Lambda, N.A. 0.75), and 60x (Nikon #MRD01605, CFI60 Plan Apochromat Lambda, N.A. 1.4) objectives, operated by NIS-Elements AR v4.5 acquisition software. Laser intensities and acquisition settings were established for individual channels and applied to entire experiments, and images were collected at the following resolution: 10x-1.73 µm/pixel, 20x-0.62 µm/pixel, 60x-0.29 µm/pixel. Image analysis was conducted using Nikon Elements, ImageJ, and Adobe Photoshop for Figure purposes. Brightness was adjusted uniformly across all pixels for a given experiment for Figure visualization purposes. Quantification of fluorescence intensities was conducted by imaging 3-5 image frames per biological replicate, which were averaged to generate a single biological replicate value. The averaged value for each replicate is depicted as open circles in each graph. Cells were selected at random while imaging surface and total receptor labeling in HEK293T cells.

#### Indirect Arrestin Assay

The indirect arrestin assay was performed using the NanoBiT membrane recruitment system reported in Spillman *et al*., 2020. HEK Arrestin 2/3 K.O cells (Alvarez-Curto *et al*., 2016) were plated into 12-well plate at a density of 3-4 × 10^5^ cells in 1 mL per well in 1x DMEM (Gibco Cat# 11995065) plus 10% FBS (Gibco Cat# 16000044) and 1X Penicillin-Streptomycin (Corning Cat# MT30002Cl). After 16-24 hours, cells were co-transfected via TransIT-2020 with receptor of interest (0.3 µg/µL); membrane-anchored AG10-CAXX LargeBit (0.05 µg/µL) and β-Arrestin-SmallBit (0.05 µg/µL); *Mm* Grk2 (0.3 µg/µL), and empty vector pEB (0.3 µg/µL) for a total of 1 µg DNA per transfection condition. Each condition required 96 µL of room-temperature 1x Opti-MEM (Gibco Cat# 31985070), 1 µL each DNA plasmid at specified concentrations), and 3 µL of room-temperature and gently-vortexed TransIT-2020 reagent. The TransIT-2020:DNA complexes mixture were gently mixed via pipetting 10 times and incubated at room temperature for 20 mins before adding drop-wise in the well. The plate was rocked gently side to side and incubated at 37°C 24 hours before harvesting. In each well, media was aspirated, and cells were washed with 1 mL warm PBS. Cells were detached with 300 µL warm Versene (Gibco Cat# 15040066) and incubated at 37°C for 7 mins then resuspended via pipetting 10 times. Cells were transferred to 1.5 mL Eppendorf tube, spun down at 500xg for 5 mins, resuspended in 1.5 mL blank 1x DMEM after supernatant were removed, and plated at 70 µL per well in Matrigel-coated 96-well assay plate. Twenty-four hours after plating, cells were treated with 20 µL 1x Nano-Glo Live Cell Substrate (Promega Cat# N2011) diluted in PBS and Arrestin assays were performed. Total luminescence of cells was measured for 0.5 sec/well in repeated manner at 42 sec cycle length for a total 1 hour. Ten µL of ligand (10^-5^ M isoproterenol or Thrombin (Sigma Cat# T4648) at 100 U/mL) or Thrombin vehicle (0.1% BSA (Roche Cat# 10735086001) in PBS) was injected at cycle 15th to reach 100 µL volume in each well.

#### Statistics

All data are expressed as means ± SEM and represent the results of at least three independent biological replicates, as indicated within each Figure Legend and as open circles within bar graphs. Statistical significance was determined using the two-tailed Student’s t-test, one-way ANOVA, or two-way ANOVA, as indicated in the Figure Legends. Data analysis and statistics were performed with Microsoft Excel, GraphPad Prism 8.0 and GraphPad Prism 9.0.

## Acknowledgements

We thank Drs. Vsevolod Gurevich and Chen Zheng (Vanderbilt University) for kindly sharing GKO HEK293 and Arr-2/3 HEK293 KO cells and for providing essential feedback, guidance, and suggestions on this study. We thank Dr. Asuka Inoue (Tohoku University) for originally providing GKO HEK293 cells. We thank Drs. Ege Kavalali, Lisa Monteggia, and all members of the Sando laboratory for critical feedback on the manuscript and study. This study was supported by grants from the NIH (R00-MH117235 to RS and R35-GM148412 to DA) and Sloan Research Fellowship (Alfred P. Sloan Foundation) to RS.

## Author Contributions

H. Bui performed all BRET2 studies, indirect arrestin assays, and immunocytochemistry/confocal experiments. A. Roach, E. Orput, and R. Raghavan performed all RNA *in situ* experiments. S. Bandekar and J. Li conducted CELSR1-3 cleavage assays. D. Araç and R. Sando designed the study with input from all the authors. R. Sando conducted all molecular cloning and wrote the manuscript with input from all authors.

