## Supplemental Online Material for "The adhesion GPCRs CELSR1-3 and LPHN3 engage G proteins via distinct activation mechanisms"

### SUPPLEMENTAL DATA FIGURES and FIGURE LEGENDS

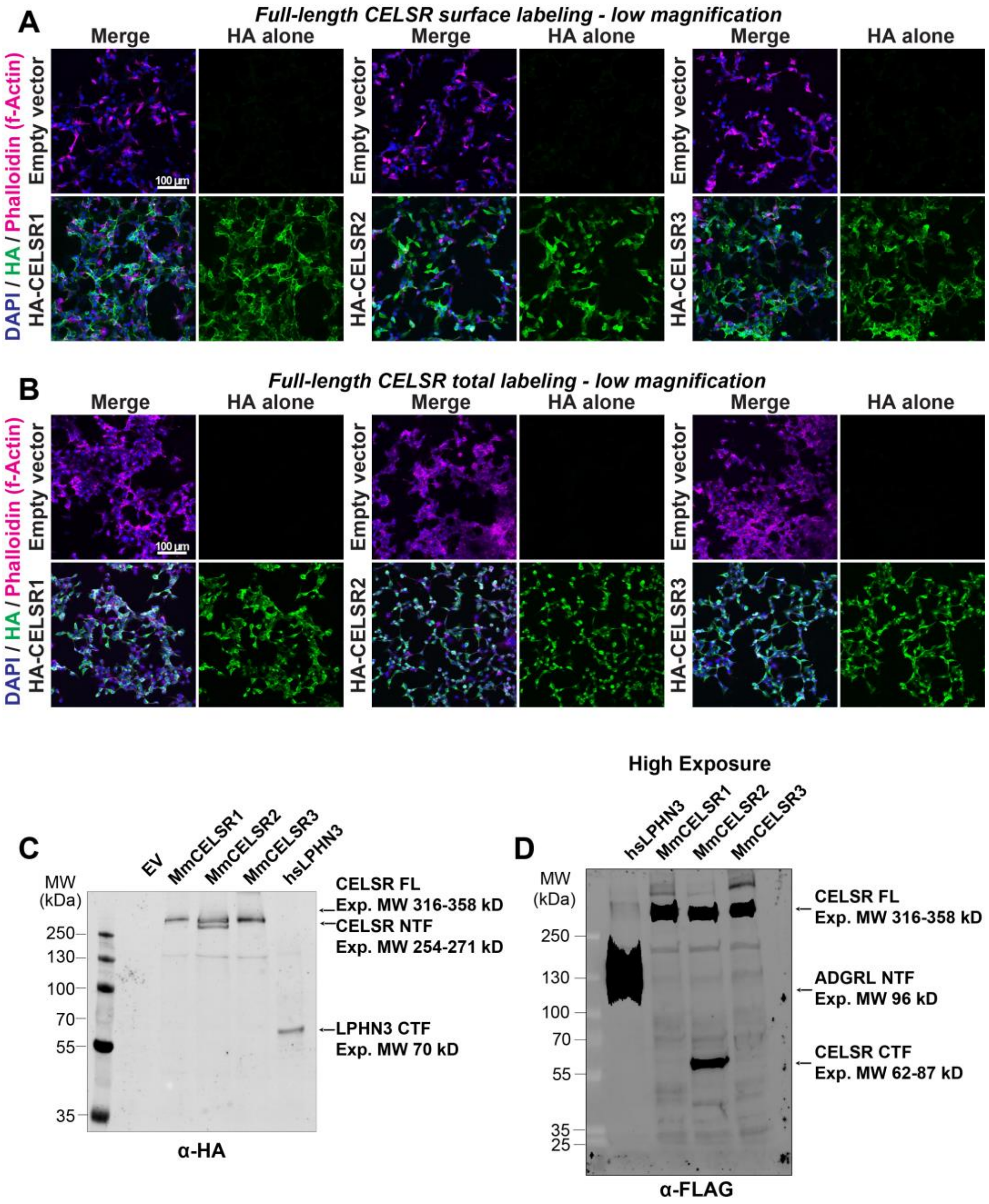


Figure S1: **Additional imaging and immunoblotting analysis from Figure 1.**

**A,** low-magnification images of HA-tagged full-length CELSR1-3 surface labeling depicting overall field-views of cell intensity distribution. Cells were transfected with indicated full-length HA-tagged *Mm* CELSR constructs or empty vector, and immunolabeled for surface HA in unpermeabilized conditions, followed by Phalloidin (f-Actin) and DAPI as internal controls.

**B,** total HA-tagged full-length CELSR1-3 labeling at low-magnification. Same as A, except cells were permeabilized with 0.2% triton X-100 prior to blocking and primary antibody incubation.

**C,** Anti-HA blot depicting detection of N-terminal cleavage product for *Mm* CELSR2 but not *Mm* CELSR1 or CELSR3. Expected sizes (kDa): *Mm* CELSR1 FL – 330; NTF – 267; CTF – 63; *Mm* CELSR2 FL – 316; NTF – 254; CTF – 62; *Mm* CELSR3 FL – 359; NTF – 271; CTF – 87; *hs* LPHN3

FL – 166; NTF – 96; CTF – 70. Note that for the *hs* LPHN3 positive control, the HA tag is present on the C-terminal fragment.

**D,** higher exposure of anti-FLAG immunoblot from Figure 1E.

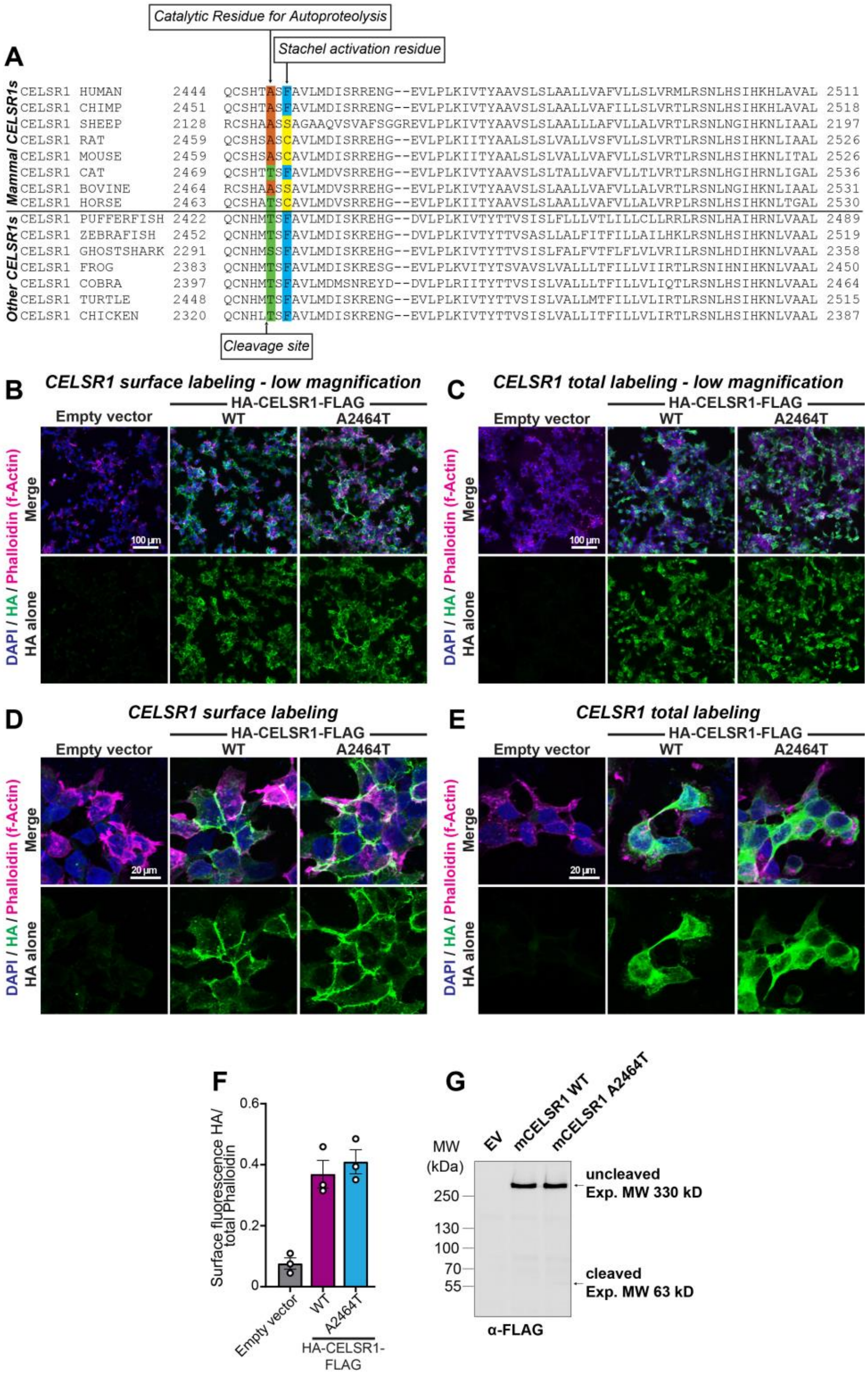


Figure S2: **Vertebrate CELSR1 sequences diverge in key residues for autoproteolysis and TA-mediated agonism.**

**A,** multiple sequence alignment of vertebrate CELSR1 orthologs. Mammalian vs. non-mammalian sequences are separated by a black horizontal line. The autoproteolytic cleavage site is shown with an arrow on the bottom of the alignment, while residues important for autoproteolysis and TA-mediated agonism are depicted on the top of the alignment.

**B,** representative low-magnification surface localization images of WT or mutant A2464T *Mm* CELSR1, which contains a catalytic threonine introduced into the location important for autoproteolysis that is present in some mammalian CELSR1 orthologs but not mouse (see panel A), compared to empty vector controls. HEK293T cells were transfected with the indicated constructs and immunolabeled for surface HA in unpermeabilized conditions, followed by labeling for Phalloidin (f-Actin) and DAPI as internal controls.

**C,** total labeling at low-magnification for indicated experimental conditions. Same as B, except cells were permeabilized with 0.2% triton X-100 prior to blocking and primary antibody incubation.

**D,** high-magnification images of cell surface labeling for HA-tagged WT CELSR1 or A2464T CELSR1 compared to empty vector controls.

**E,** total pool of HA-tagged receptors in indicated experimental conditions.

**F,** quantification of HA surface intensity from high-magnification images relative to Phalloidin from indicated experimental conditions. Total cell fluorescence was measured for HA and Phalloidin channels from three independent biological replicates.

**G,** *Mm* CELSR1 autoproteolysis assay. HEK293T cells were transfected with the indicated HA (N-terminal) and FLAG (C-terminal) tagged CELSR1 constructs of empty vector control and subsequently immunoblotted for the C-terminal FLAG to assess autoproteolysis. Representative blot depicts that neither WT nor mutant A2464T CELSR1 exhibits predominant cleavage. The C-terminal cleavage product is estimated to be 63 kDa, while the full-length uncleaved receptor is 330 kDa.

Numerical data are means ± SEM from 3 independent biological replicates (depicted as open circles).

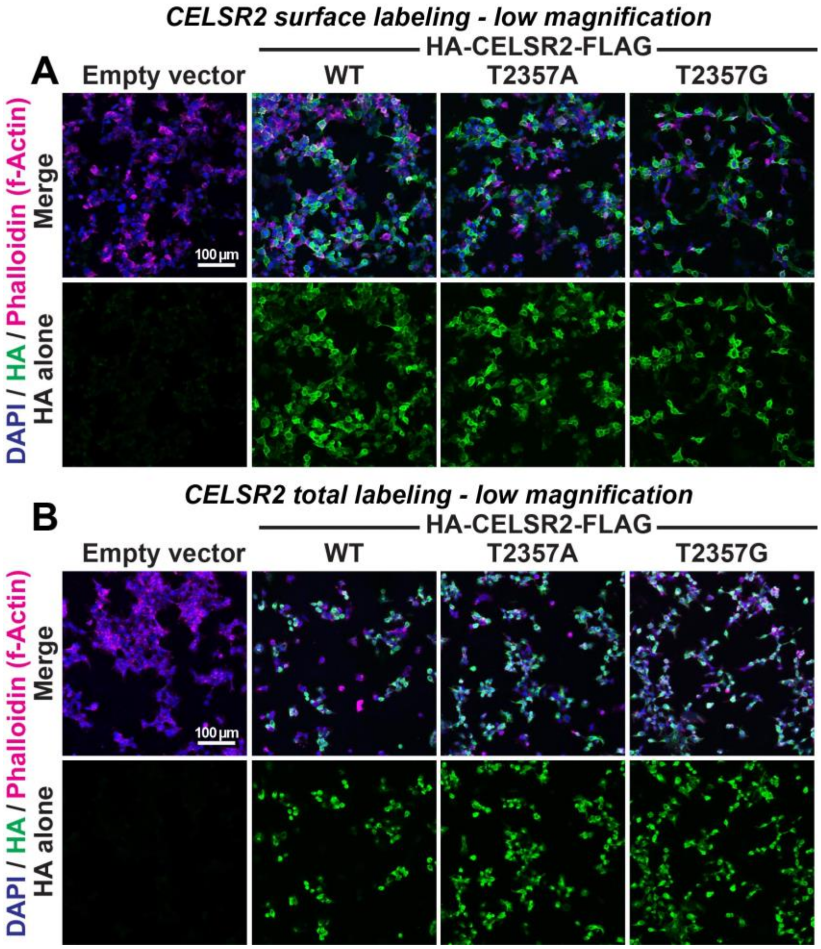


Figure S3: **Low-magnification images of full-length WT, T2357A, and T2357G CELSR2.**

**A,** representative low-magnification images of surface labeling for overexpressed HA-tagged WT, T2357A, and T2357G CELSR2 compared to empty vector controls. HEK293T cells were transfected with the indicated constructs and immunolabeled for surface HA in unpermeabilized conditions, followed by labeling for Phalloidin (f-Actin) and DAPI as internal controls.

**B,** representative low-magnification images of total labeling for indicated conditions. Same as A, except cells were permeabilized with 0.2% triton X-100 prior to blocking and primary antibody incubation to expose the total pool of receptors.

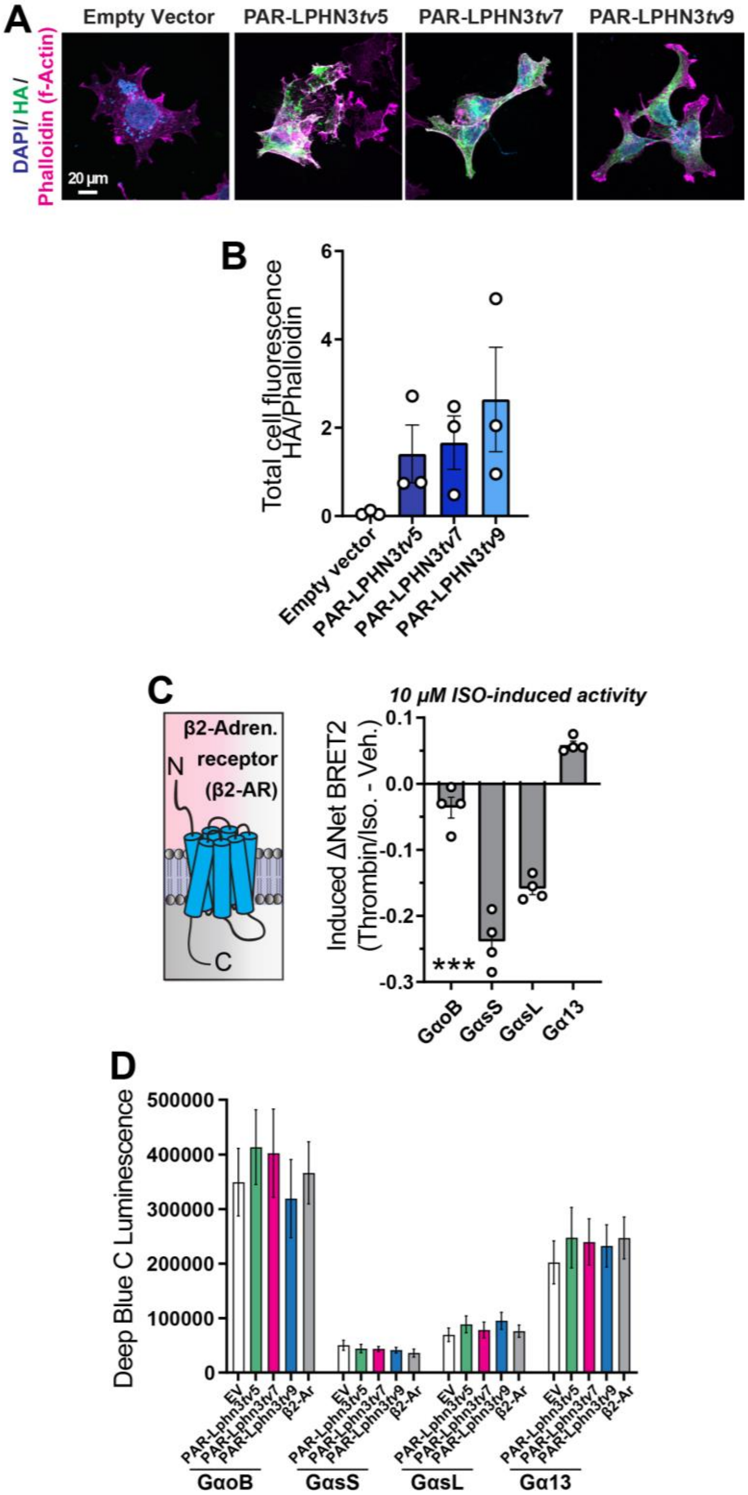


Figure S4: **Expression of PAR-LPHN3 variants in HEK293T cells and additional BRET2 data.**

**A,** Representative images of C-terminally HA-tagged PAR-LPHN3*tv*5, PAR-LPHN3*tv*7, and PAR-LPHN3*tv*9. HEK293T cells were transfected with indicated constructs or empty vector, and immunolabeled 48-hours post-transfection for HA together with Phalloidin (f-Actin) and DAPI (nuclei) as internal controls. Versions lacking the C-terminal HA tag were used for all BRET2 experiments to exclude a potential impact of the tag on G protein coupling efficacy.

**B,** quantification of HA channel intensity relative to Phalloidin. Total cell fluorescence was measured for HA and Phalloidin channels from three independent culture replicates.

**C,** Isoproterenol-induced (ISO) BRET2 with β2-adrenergic receptor (β2-AR) and indicated TRUPATH sensors. GKO HEK293 cells were transfected with the indicated BRET2 G protein sensor combinations and β2-AR, and net BRET2 signals of 10 µM agonist ISO were compared to vehicle treated cells.

**D,** Deep Blue C donor luminescence emission measured during BRET2 experiments from Figure 3. Data are from 4 independent biological replicates.

Numerical data are means ± SEM from 3-4 independent biological replicates (depicted as open circles). Statistical significance was assessed by one-way ANOVA.

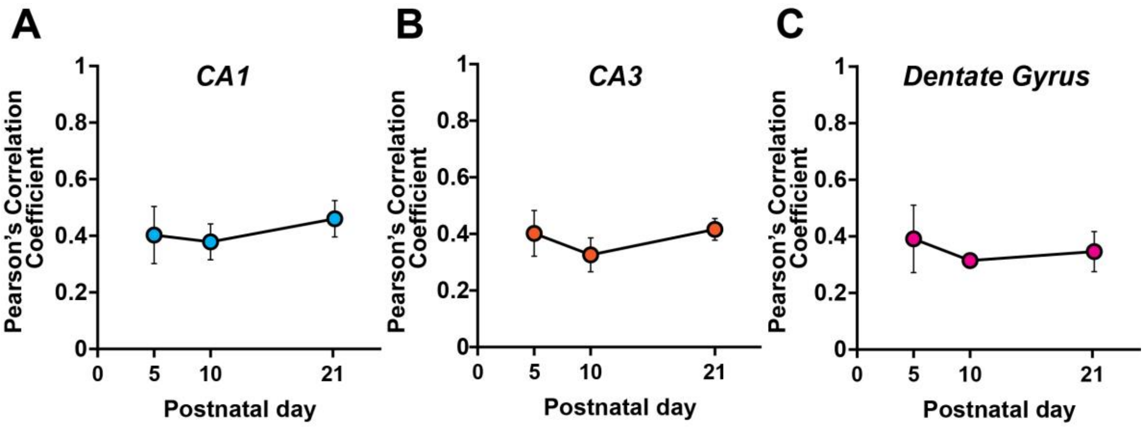


Figure S5: **RNA *in situ* hybridization analyses of *Lphn3* in the developing mouse hippocampus.**

**A-C,** quantification of the Pearson's correlation coefficient of the *pan-Lphn3* channel (green) to the *Ct-Lphn3* variant channel (red) in the CA1 (A), CA3 (B), or Dentate Gyrus (C). Quantifications are from experiments described in Figure 4.

Numerical data are means ± SEM from 3-4 independent mouse replicates.

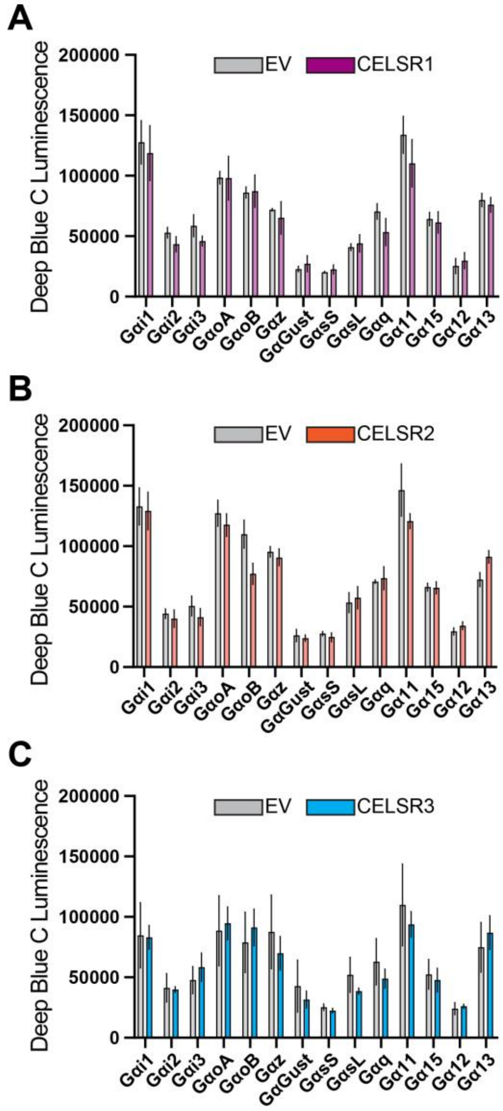


Figure S6: **Donor luminescence measurements from BRET2 experiments in Figure 5.**

**A-C,** Deep Blue C (410 nm) luminescence data from BRET2 experiments conducted in Figure 5A (CELSR1), D (CELSR2), or G (CELSR3).

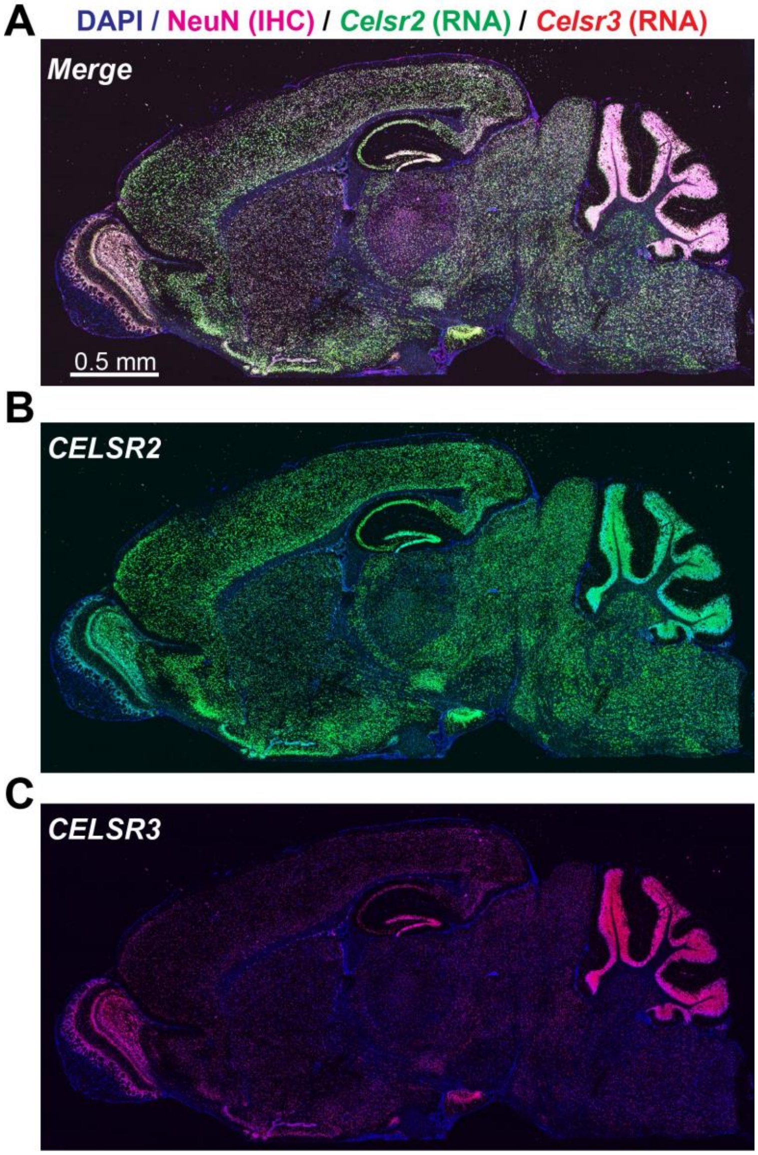


Figure S7: **Overview of *Celsr2/3* spatial expression throughout sagittal sections of the postnatal day 21 mouse brain.**

**A,** representative postnatal day 21 sagittal section co-labeled via double IHC/RNA *in situ* for the neuronal marker NeuN (IHC; magenta), *Celsr2* (RNA *in situ*, green), *Celsr3* (RNA *in situ*; red) and DAPI.

**B,** image depicting *Celsr2* expression together with DAPI from A. Note broad expression of *Celsr2* throughout the mouse brain.

**C,** same as B, except for *Celsr3*. *Celsr3* displays a more discrete spatial expression pattern compared to *Celsr2*, and is particularly enriched in regions like the olfactory bulb, hippocampus, and cerebellum, where it overlaps with *Celsr2*.

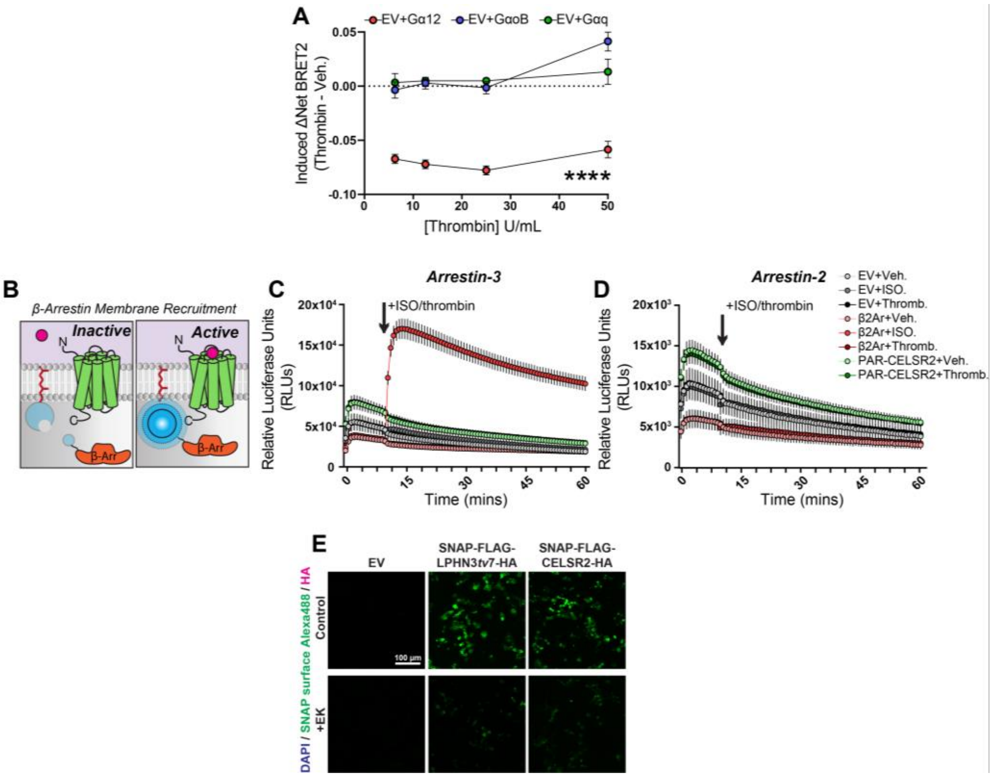


Figure S8: **Additional analyses of PAR-CELSR2 via G protein or β-Arrestin coupling.**

**A,** BRET2 analysis of GKO HEK293 cells co-transfected with empty vector and TRUPATH BRET2 combinations for Gα12, GαoB, Gαq and treated with increasing concentrations of thrombin. High concentrations of thrombin activate the Gα12 TRUPATH BRET2 sensors in the absence of exogenously expressed receptor. Data represent the mean ± SEM of 7 independent biological replicates.

**B,** Schematic diagram of Arrestin NanoBiT assays to assess Arrestin-2/3 membrane recruitment following GPCR activation. *Left*, Arrestin fused to SmBiT resides predominately in the cytoplasm in unstimulated conditions, where it is unable to interact with membrane-tethered LgBiT. *Right*, activation of overexpressed GPCR recruits Arrestin to the membrane, allowing NanoBiT reconstitution which thereby increases luminescence.

**C,** Arrestin-3 NanoBiT recruitment assay. HEK293 Arr2/3 KO cells were transfected with indicated receptors/empty vector together with membrane-tethered LgBiT, SmBiT-*Mm*Arr3, and GRK2. Baseline luminescence was measured for 10 minutes, followed by treatment with 10 µM isoproterenol (ISO) or 10 U/mL thrombin and subsequent luminescence measurements for a total of 60 minutes. The $\beta_2$Ar stimulated with ISO was used as a positive control. Data are from 3 independent biological replicates.

**D,** NanoBiT assays for indirect Arrestin-2 membrane recruitment in HEK293 Arr2/3 KO cells. Cells were co-transfected with indicated receptors/empty vector, membrane-bound LgBiT, and *Mm* Arrestin-2 fused SmBiT and *Mm* GRK2, and treated with either 10 µM isoproterenol or 10 U/mL thrombin after measuring baseline luminescence for 10 mins. $\beta_2$Ar, used as a positive control, has been shown to specifically interact with Arrestin-3 but not Arrestin-2. Data are from 3 independent biological replicates.

**E,** Isolated SNAP-surface Alexa Fluor 488 channel from images in Figure 7E.

Numerical data are means ± SEM from 3-7 independent biological replicates as indicated in the Figure Legend. Statistical significance was assessed by two-way ANOVA (****, p<0.0001).

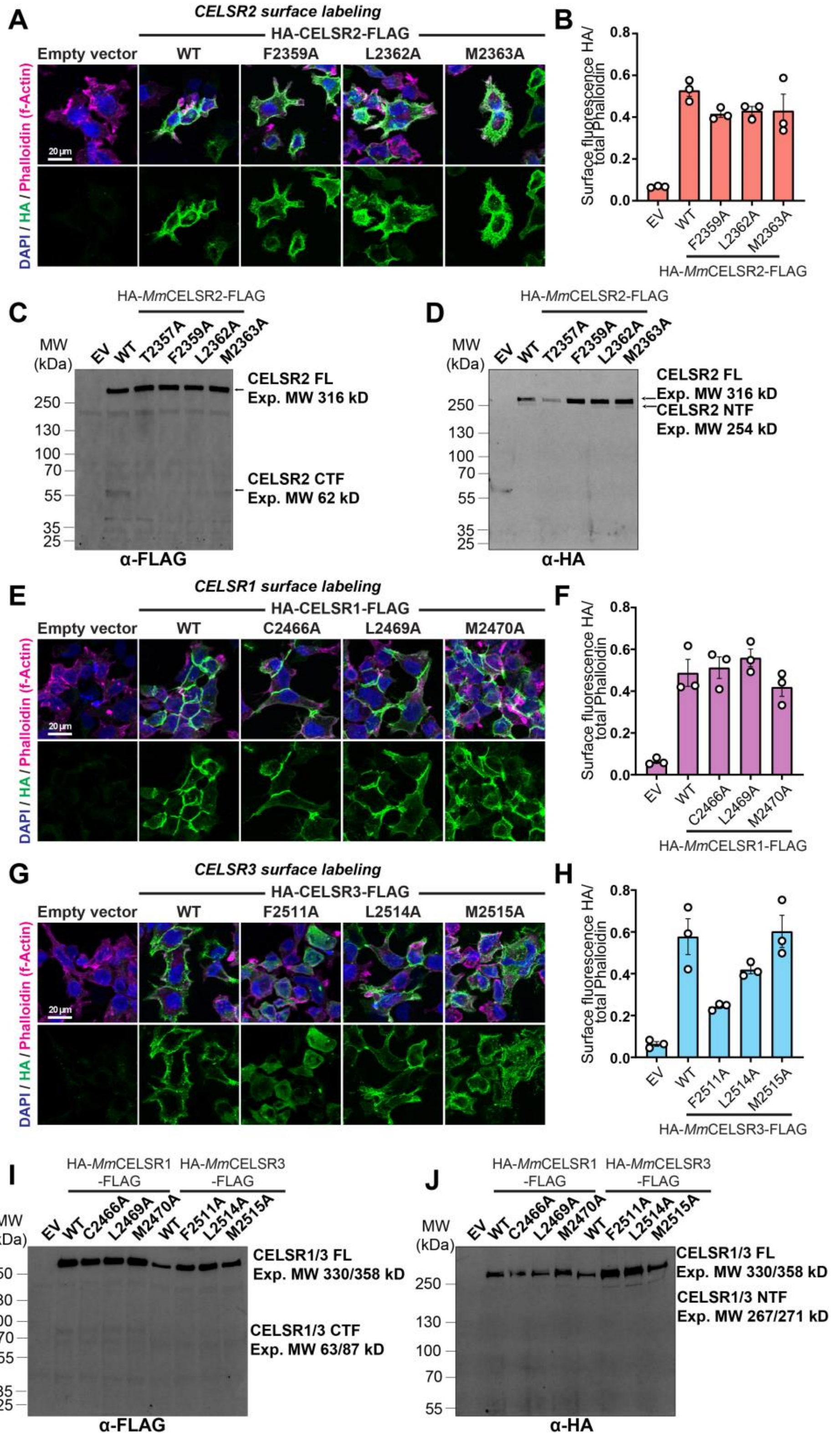


Figure S9: **Surface expression and autoproteolysis of full-length CELSR1-3 TA point mutants.**

**A,** surface expression and localization of indicated dual HA/FLAG-tagged, full-length *Mm* CELSR2 mutants compared to empty vector and WT CELSR2. Transfected HEK293T cells were labeled for surface HA in unpermeabilized cells, followed by DAPI and Phalloidin (f-Actin).

**B,** Relative HA surface intensity compared to the intensity of Phalloidin, used as an internal control.

**C & D,** full-length *Mm* CELSR2 cleavage assays in HEK293T cells. Constructs contained an N-terminal HA following a preprotrypsin signal peptide, and C-terminal FLAG tag. HEK293T cell lysates were probed for FLAG (**C**) or HA (**D**) compared to WT CELSR2 and T2357A CELSR2.

**E & F,** same as A and B, except for full-length N-terminal HA tagged, C-terminal FLAG tagged WT, C2466A, L2469A, and M2470A CELSR1.

**G & H,** Same as A and B, except for full-length N-terminal HA tagged, C-terminal FLAG tagged WT, F2511A, L2514A, and M2515A CELSR3.

**I & J,** full-length *Mm* CELSR1 and CELSR3 cleavage assays in HEK293T cells. Immunoblots were conducted as in C and D, except for indicated CELSR1 or CELSR3 mutants and WT controls. Blots were probed for C-terminal FLAG (**I**) or N-terminal HA (**J**).

Data in panels B, F, and H depict quantifications of means ± SEM from three independent biological replicates.
